# Real-time dynamics of individual chemoreceptor mRNA molecules reveals translation hotspots at the inner membrane of *Escherichia coli*

**DOI:** 10.1101/2022.12.16.520495

**Authors:** Tobias Bergmiller, Ekaterina Krasnopeeva, Srdjan Sarikas, Nela Nikolic, Calin C. Guet

## Abstract

Since bacteria lack a nucleus, the location of mRNA molecules is determined by the different characteristics of the encoded proteins, and the transcriptome is spatially arranged into cytosolic and membrane-associated mRNA. While translation of membrane protein-encoding mRNA has been studied in great mechanistic detail using biochemical methods, the spatiotemporal dynamics of this process remains poorly understood at the subcellular level. Here, we investigate the dynamics of individual fluorescently labelled mRNA molecules encoding the transmembrane serine chemoreceptor Tsr, to probe the mechanism of membrane protein translation. Analysis of *tsr* mRNA diffusion in the proximity of the plasma membrane revealed distinct diffusive modes that reflect the state of the mRNA molecule and its involvement in the process of active translation into the Sec secretion system. We find that the composition, and hence the fluidity of the membrane affects diffusion of membrane targeted mRNAs. Moreover, Tsr translation occurs in localized membrane regions, similar to eukaryotic hotspots. The hotspot localization coincides with the physical location of the transcribed gene, which itself is displaced towards the inner membrane. These findings suggest that inner membrane protein translation is a spatially defined process that occurs in hotspots, indicative of long-lived transertion sites. Our results show an additional layer of spatio-temporal structuring within the bacterial cell, thus revealing a qualitatively different understanding of the basic process of transcription and translation in bacteria.

**Significance Statement:** A large fraction of the bacterial proteome is directly synthesized into the inner membrane, and this process shapes the overall distribution of mRNA transcripts within the cell. Although highly dynamic in their nature, bacterial transcriptomes have mostly been studied in fixed cells. Here, we track individual mRNA molecules encoding the serine chemoreceptor in living bacterial cells and find that translation occurs in membrane hotspots that were previously exclusive to eukaryotes. Our results indicate an additional layer of spatio-temporal structuring within the bacterial cell that impacts our understanding of transcription and translation in bacteria.

## Introduction

Bacteria are highly structured entities at the subcellular level. The nucleoid structure and the distribution of RNA polymerases and ribosomes within the cell are highly organized and orchestrated biological processes (1-4). The surrounding inner membrane and outer membrane appear to have an intricate structure (5) (6)characterized by functional microdomains (7). However, the spatial distribution and organization of the bacterial transcriptome, is much more dynamic compared to many macromolecular building blocks of the cell, and therefore much harder to experimentally capture.

Different models of transcriptome organization across bacterial species have been proposed (8). In *Caulobacter crescentus* specific mRNAs and their corresponding proteins were shown to be localized in the vicinity of their origin of expression (9). Several reports describe overall mRNA distribution according to ribosome density that peaks at the outer edges of the chromosome (10) (2), suggesting global mRNA localization mechanisms that rely on both, exclusion of mRNA from the densely packed chromosome and translation-driven localization. However, a growing body of work indicates translation-independent mRNA and small regulatory RNA (sRNA) localization, where dynamic and stress-dependent localization patterns have been observed (11, 12). In these models, mRNA and sRNA distribution and localization are defined by the interaction with regulatory hubs such as Hfq rather than ribosomes (12).

mRNA targeting to the Sec translocation machinery has recently been identified as a key factor determining the organization of the transcriptome and in particular transmembrane-encoding mRNAs in *E. coli* (13). While mRNA molecules encoding periplasmic, outer-membrane and cytosolic proteins are predominantly localized in the cytosol, mRNA species encoding transmembrane proteins are sequestered to the inner membrane by co-translational targeting through the signal recognition particle (SRP) pathway (14). About 20% of all *E. coli* proteins are co-translationally inserted into the inner membrane through the Sec translocon (**Figure 1A** (15): after translation of a trans-membrane or integral membrane protein is initiated, SRP engages on hydrophobic nascent chains emerging from the ribosome exit tunnel. SRP then “sorts” the ribosome-nascent chain complex (RNC) with bound mRNA through interaction with the receptor FtsY to the inner membrane (16). FtsY facilitates 2-dimensional diffusion of the complex along the inner membrane until the RNC binds to SecY at the SecYEG translocon (17)(**Figure 1A**). Consequently, membrane targeted RNCs and associated mRNAs localize to the inner membrane, leading to compartmentalization of a subset of mRNA species through translational targeting (13). Furthermore, the process of Sec-dependent protein expression is subject to transertion, which is the process of close coupling of transcription, translation, and Sec-dependent insertion in bacteria (18). Transertion involves displacement of genomic loci encoding transmembrane proteins towards the membrane and was postulated to contribute to the formation of membrane inhomogeneities and hyperstructures, which are dynamic assemblies of interacting macromolecules (19).

**Figure 1:**
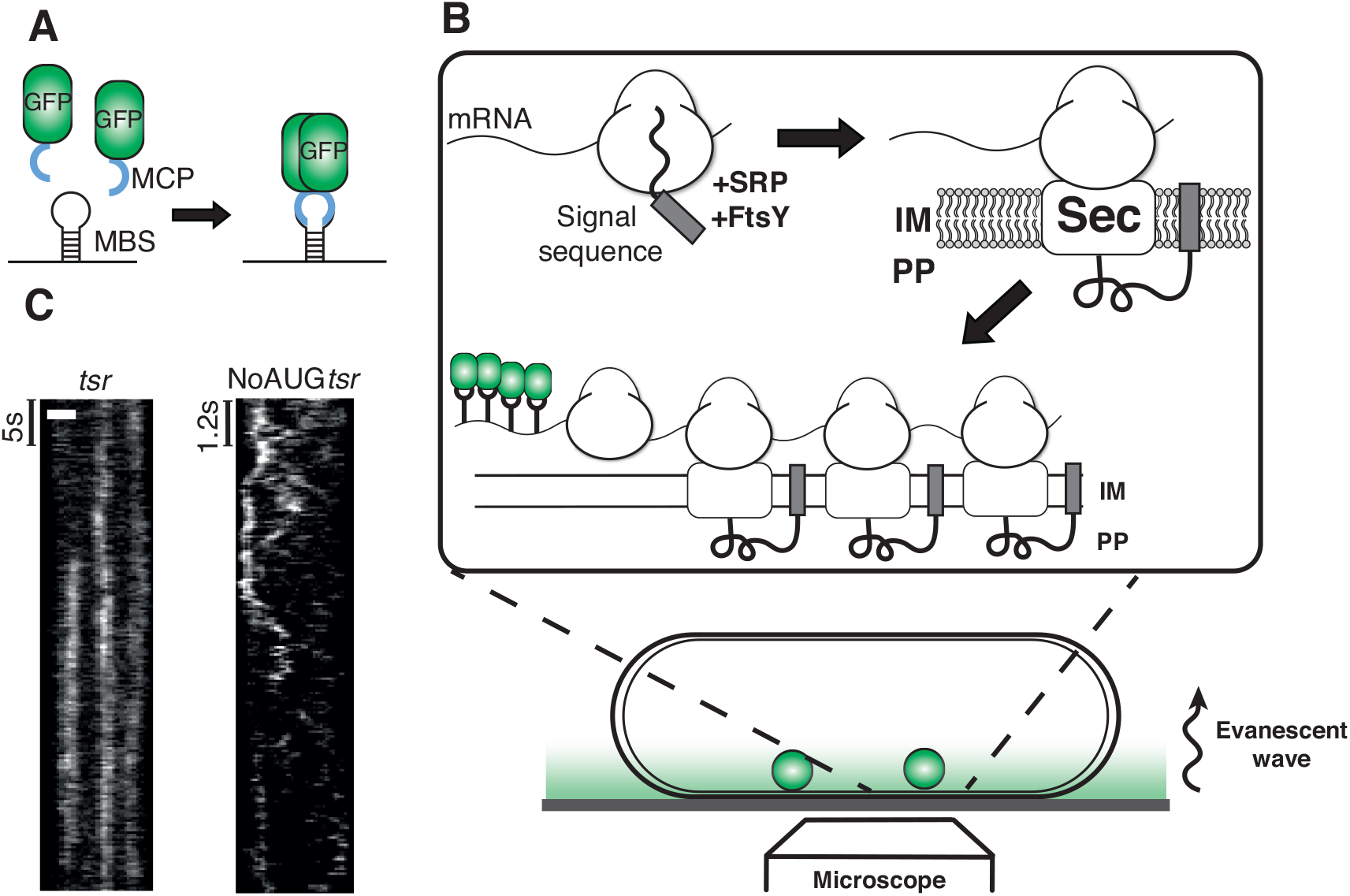
MS2 fluorescent labelling of *tsr* mRNA allows to track individual mRNA molecules in living bacteria. a) Proteins encoding a strongly hydrophobic N-terminus, such as the chemoreceptor Tsr, are translated through the co-translational insertion pathway. Exposure of a hydrophobic N-terminus, serving as signal sequence, at the ribosome exit tunnel after initiation of translation leads to engagement of signal recognition particle (SRP) and the SRP receptor protein FtsY. Upon membrane contact, the mRNA-ribosome-RNC complex interacts with the Sec translocon, so that Tsr is translated directly into Sec and the membrane through the interaction between ribosome and Sec. Long transcripts are then bound by multiple ribosomes that engage with several Sec translocons, forming a membrane-bound polysome. b) The MS2 system consist of two components derived from MS2 phage: a MS2 capsid protein (MCP) and a MS2 binding site (MBS) forming a secondary RNA hairpin structure. MCP binds as a dimer to a single mRNA containing a MBS, and this interaction can be visualized using fluorescence microscopy when MCP forms a protein fusion with GFP. c) Kymograph representations of *tsr* dynamics in individual cells. Left: *tsr* displays longitudinal patterns along the cell axis. Right: fast noAUG-*tsr* dynamics imaged using illumination with a highly oblique angle (HILO TIRF). Scale bar = 1*μ*m

Despite a growing understanding of spatial transcriptome organization in bacteria, many questions remain open. The spatiotemporal dynamics of individual mRNA molecules that undergo translation through transertion remains poorly understood. It is also unclear how the physical properties of mRNA molecules such as size or the fluidity of the bacterial membrane affect this process. Are translation sites of individual mRNA molecules distributed uniformly along the inner membrane or organized into localized “translation hotspots”? Such hotspots, displaying high densities of actively translated mRNA due to their confinement in ribosome-enriched areas, have been reported in eukaryotes (20) but not in prokaryotes.

With few exceptions (21, 22), the majority of recent studies is limited by the choice of experimental methods that do not capture the dynamics of individual mRNA molecules and cannot distinguish between actively translated and unbound mRNA. Here, we study the diffusive dynamics and the spatial distribution of individual mRNA molecules in living *Escherichia coli* cells using the serine chemoreceptor Tsr as an example transmembrane protein that is translated directly into the membrane by the Sec translocation machinery (23).

## Results

### Tracking individual *tsr* mRNA molecules in real time in living *Escherichia coli* cells

To image individual mRNA molecules in living cells, we used the MS2 labelling system pioneered by the Singer lab (24), which consists of two components: a fluorescently labelled RNA phage MS2 coat protein (MCP), and a secondary structure RNA element forming a unique hairpin (MS2 binding site, MBS, **Figure 1A**) (25), which can be encoded as part of any mRNA molecule. The MBS is recognized by the MCP with high specificity and affinity. We used a system similar to the one used in (26), with the difference that the monomeric delFG mutant MCP was modified with a high-affinity mutation (N55K (27)) and fused to a monomeric superfolder GFP (sfGFP) (28). We refer to this fusion as MS2-sfGFP hereafter. To maximize the signal-to-noise ratio, the MS2-sfGFP fusion was expressed under control of the P_LlacO1_ promoter at low levels (5 *μ*M IPTG) from a very low-copy plasmid (SC101* origin of replication (29)) in the presence of high levels of LacI repressor (for strain details see **Material and Methods**). This resulted in reduced fluorescent background due to low amounts of unbound MS2-sfGPF and no detectable fluorescent spots in the absence of MBS-containing mRNA (**Supplementary Movie 1**).

We tracked mRNA particles encoding Tsr chemoreceptor by placing 24 MBS repeats onto the 3’-untranslated region (3’-UTR) of the *tsr* mRNA, which did not interfere with expression of Tsr receptors. The latter was verified by fusing Tsr to photoactivation mCherry (PAmCherry) and visualizing Tsr-PAmCherry protein expression (**Supplementary Movie 2**). To control transcription of *tsr* mRNA, we used the inducible and tightly repressed P_LtetO1_ promoter controlled by TetR (29), and induced *tsr-24xms2* expression using sub-saturating amounts of the inducer anhydrous tetracycline (aTc, 5ng/ml). While tight MS2-sfGFP*-*MBS binding hinders mRNA degradation (30), our focus was on maximizing the signal-to-noise ratio for tracking mRNA particles over periods of up to several minutes, instead of capturing the full mRNA life cycle.

In order to image *tsr* mRNA diffusion and membrane targeting at the inner membrane (**Figure 1B**), we used TIRF microscopy. Co-expression of MS2-sfGFP and *tsr-24xms2* (abbreviated *tsr* hereafter) yielded bright fluorescent spots in near proximity of the inner membrane (**Figure 1C, Supplementary Movie 3**). Kymograph representations of individual cells showed at least two distinct modes of *tsr* mRNA particles behavior: transient and prolonged particle walks at the membrane creating straight vertical traces of *tsr* molecules, and rapid movement with complex trajectories for non-translating noAUG*tsr* molecules. Additional smFISH labelling in the absence of MS2-sfGFP ruled out MS2-sfGFP-induced localization artifacts and confirmed characteristic patterns of *tsr* fluorescent spots at the bacterial membrane in the presence of the transcriptional inducer aTc (**Supplementary Figure 1**).

### Diffusive behavior of *tsr* mRNA reveals distinct translation-dependent subpopulations

To gain quantitative insights into the diffusive behavior of *tsr* mRNA, we tracked single particles and performed mean square displacements (MSD) analysis (for details see **Material and Methods**). The distribution of the apparent 2D diffusion coefficients calculated from the MSD for one time interval is shown in **Supplementary Figure 2**. Our fitting procedure showed that *tsr* mRNA displayed three distinct diffusive states with diffusion coefficients ranging approximately from 0.001 *μ*m^2^/s for near-stationary slow-moving particles, to 0.05 *μ*m^2^/s for rapidly moving molecules, irrespective of whether the mRNA molecule was expressed from a plasmid or from a single-copy insertion into the chromosome. The results of this fitting procedure and all fits to other mRNA variants are presented in **Figure 2** and **Supplementary Figure 2 and Supplementary Figure 3**. This is in good agreement with a recent finding that mRNA diffusion in *Bacillus subtilis* shows up to three diffusive states (21), although our bacteria of choice, labelling system and imaging conditions were overall different. A construct with only 6 MBS elements, *tsr-6xms2*, showed a nearly identical distribution of subpopulations (**Figure 2A**).

**Figure 2:**
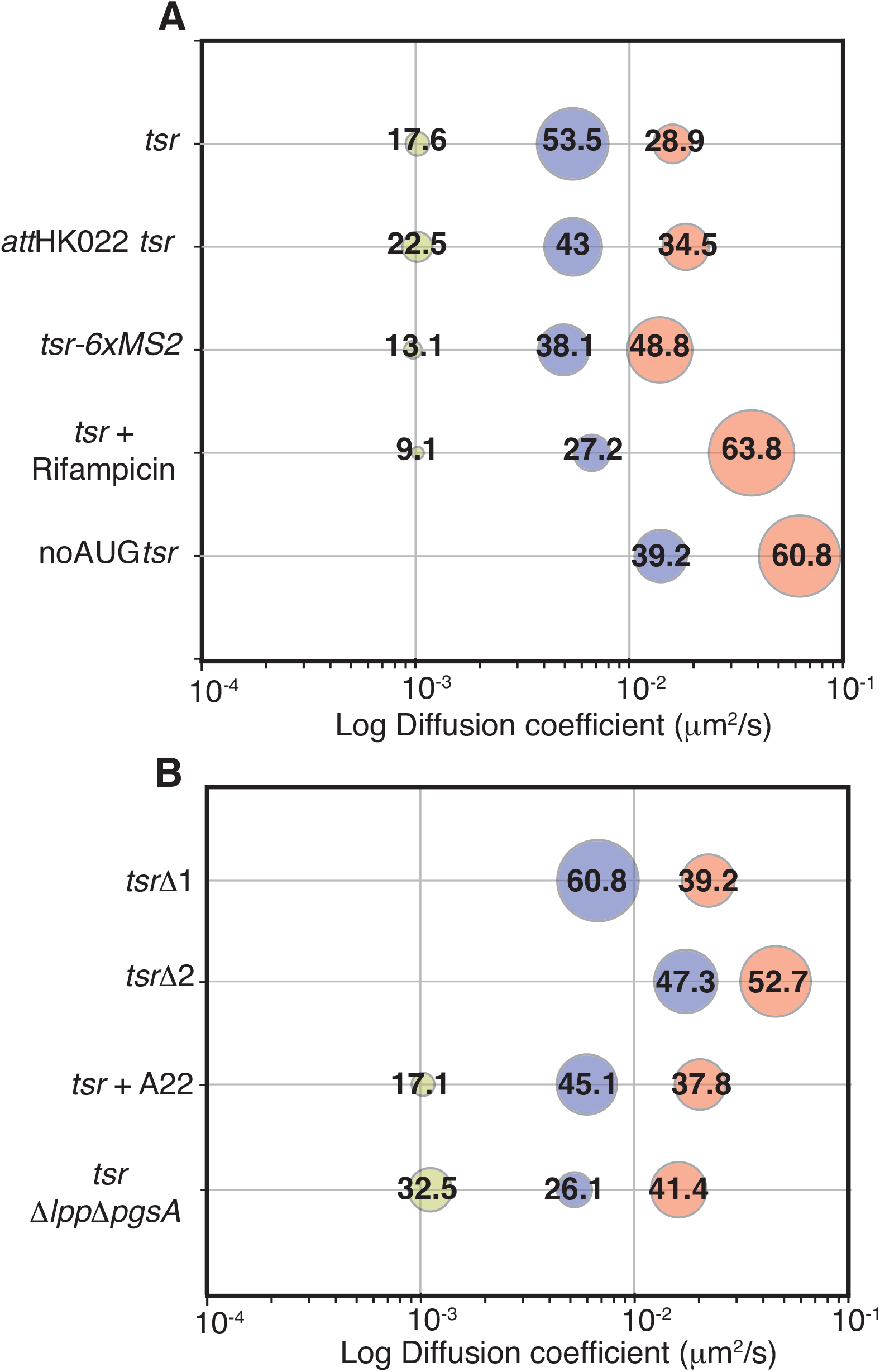
*tsr* mRNA molecules can be classified into subpopulation according to their diffusive behavior, and diffusion changes with mRNA length and membrane perturbation. a) Bubble plot showing the summary of the fitting procedure for different mRNA species. Numbers on top of bubbles and bubble size indicate the weight of each subpopulation. X-axis: log diffusion coefficient, *μ*m^2^/s. *tsr*: mRNA encoding Tsr, labelled with 24 MBS and expressed from a pZA21 plasmid. *tsr (att*HK022) – same as *tsr* but expressed from the *att*HK022 site of the chromosome. *tsr-6xms2 –* mRNA encoding Tsr protein, labelled with 6 MBS and expressed from a pZA21 plasmid; *tsr +*Rifampicin – 25 *μ*g/ml Rifampicin added to cells expressing *tsr*. noAUG-*tsr– tsr* with the start codon removed. b) *tsr*Δ*1– tsr* with the removed cytoplasmic coiled-coil domain ΔAA 215-551; *tsr*Δ*2 – tsr* truncated to have the first transmembrane α-helix only, ΔAA 31-551; *tsr+*A22 *–* 10 *μ*g/ml A22, causing depolymerization of MreB, added to the cells expressing *tsr. tsr* Δ*lpp*Δ*pgsA– tsr* is expressed in the strain lacking phosphatidylglycerol and cardiolipin.

To characterize the three diffusive states of *tsr* mRNA further, we expressed and tracked two different species of mRNA molecules that could not be translated: a noAUG*tsr* mRNA, lacking the start codon, and an mRNA consisting of 24 MBS only. We found that the corresponding single-step distributions were devoid of the subpopulation with the lowest *D*, while the subpopulation of fast-moving particles was overrepresented (**Figure 2A and Supplementary Figure 3**). This is in good agreement with the finding that active translation by ribosomes shields mRNAs from degradation (31), suggesting that fast moving particles are most likely degradation products. To further test this assumption, we treated cells expressing *tsr* with rifampicin, an inhibitor of RNA polymerase (RNAP). Rifampicin binding to RNAP causes sudden transcriptional arrest, and subsequently the transcription-degradation equilibrium of the cellular mRNA pool is shifted towards degradation (32). The results showed a strong shift in the subpopulation frequencies towards particles with high *D*, and a substantial reduction of particles with low *D*, in support of fast-moving particles representing degradation products (**Figure 2A)**.

We hypothesized that the slow-diffusing mRNA subpopulation is characteristic for co-translationally targeted transcripts that undergo active translation at the inner membrane. To test this, we imaged *lacY-24xms2*, and *cfp-24xms2* mRNAs, which encode a different transmembrane protein, lactose permease LacY, and the cytosolic cyan fluorescent protein (CFP), respectively. In agreement with our assumption, *cfp* displayed only the intermediate and fast subpopulation, while *lacY* showed a third low-*D* subpopulation (**Supplementary Figure 3**).

According to the model of co-translational protein insertion of Tsr into the membrane via the Sec translocon, and assuming an average ribosome occupancy of 1.3 ribosomes per 100 nucleotides (33), the *tsr* mRNA is tethered to the bacterial inner membrane through binding of about 21 ribosomes to SecYEG secretion systems in the fully saturated state. This process of membrane polysome formation explains our observation of near-stationary diffusive subpopulations in membrane-targeted mRNAs that is a magnitude lower than the diffusion constant of large transmembrane proteins with more than a dozen transmembrane helixes (34).

We further assumed that the intermediate subpopulation consists of mRNA particles bound to ribosomes but not tethered to the inner membrane, since all transcripts, irrespective of the presence or absence of a coding region or a start codon, carry a very strong ribosome binding site (RBS) at the 5’ UTR. The RBS is likely to attract binding of a pre-initiation complex consisting of the 30S ribosomal subunit and additional factors such as fMet-tRNA (35), subsequently substantially increasing the molecular mass, which affects the apparent *D* of the molecule. Of note, the intermediate subpopulation matches the previously published apparent *D* of actively translating ribosomes (2).

We concluded that the three modes of diffusive behavior can be approximated as: (i) slow and Sec-translocon-associated translating particles (diffusion constant *D* of approximately 0.001 *μ*m^2^/s), (ii) intermediate fraction of ribosome-bound particles (*D* of 0.005 to 0.01 *μ*m^2^/s), and (iii) fast particles consisting of free mRNAs or degradation products with a *D* ranging from 0.015 to 0.05 *μ*m^2^/s.

### Transcript length and membrane fluidity affect *tsr* mRNA diffusive behavior

To gain a better understanding of the diffusive behavior, we set out to identify some of the factors affecting *tsr* mRNA diffusion. We first investigated whether the frequency of low-*D* particles depends on mRNA length, which determines the predicted polysome size as well as the extent of tethering to the inner membrane. The Tsr protein consists of two transmembrane α-helices flanking a large periplasmic domain, and a long C-terminal cytosolic multimerization domain (36). We built several Tsr mutants by removing the cytoplasmic coiled-coil domain (ΔAA 215-551, *tsr*Δ*1* hereafter), or by expressing the first transmembrane α-helix only (ΔAA 31-551, *tsr*Δ*2* hereafter), thus reducing the assumed average ribosome occupancy from approximately 21 to about 10 ribosomes and 1-2 ribosomes per transcript, respectively. Our analysis showed that variant *tsr*Δ*1* yielded a subpopulation distribution lacking the low *D* subpopulation (**Figure 2B**). Thus, shortening *tsr* mRNA by around 60% substantially reduced tethering of the transcript to the membrane. *tsr*Δ*2*, in turn, not only lacked the low-*D* subpopulation, but also showed an overall increase in the fast population and a shift of both subpopulations towards higher *D*, supporting our hypothesis that a short mRNA is either insufficiently or very transiently membrane-targeted due to low ribosome occupancy or small nascent peptide size (37). Furthermore, the short transcript *tsr*Δ*2* encodes a 30AA peptide that is likely too poor of a signal for sorting the RNC-mRNA complex to the Sec translocon, as the completed TsrΔ2 will emerge from the ribosome exit tunnel only after translation is being terminated.

The difference in molecular mass between the different mRNA variants cannot account for the differences we observed in our experiments. For instance, a full-length *tsr-24xms2* mRNA molecule saturated with 48 MS2-sfGFP proteins has a molecular mass of approximately 2.3 MDa, which is below the molecular mass of a single fully assembled bacterial ribosome of 2.7 MDa (38). This discrepancy between molecular mass of fully decorated mRNA particles and hypothetical polysome sizes agrees with our finding that the total size of decorated MBS repeats (24 versus 6 repeats) play a minor role in the observed differences in diffusion. A full comparison between different mRNA lengths, predicted polysome size and molecular mass is listed in **Supplementary Table 1**.

We next tested whether membrane fluidity contributes to the diffusive behavior of membrane-targeted mRNAs. We hypothesized that a decrease in membrane fluidity by removal of anionic phospholipids (APLs) or depolymerization of MreB, will reduce *tsr* mRNA diffusion similar to how it affects other membrane-associated processes, such as cytokinesis, envelope expansion, and maintenance of membrane potential (39). Phosphatidylglycerol (PG) and cardiolipin (CL) are anionic phospholipids of the *E. coli* membrane, and CL is localized to the septum and cell poles (40). Low levels of PG and CL reduce membrane fluidity (41) and absence of CL destabilizes SecYEG translocon stability and integrity (42). We measured *tsr* diffusion in a Δ*lpp*Δ*pgsA* strain TB334, which is largely devoid of PG and CL (43). TB334 displayed severe defects in morphology and growth (**Supplementary Movie 4**). Importantly, *tsr* mRNA displayed an increased frequency of the low *D* subpopulation (from ∼20 to above 30%), supporting our hypothesis that decreasing membrane fluidity slows down *tsr* mRNA diffusion (**Figure 2B**).

MreB is an actin-like cytoskeletal protein that moves circumferentially along the inner membrane of *E. coli*, and its motion is coupled to peptidoglycan synthesis (44). It has been shown that MreB affects membrane dynamics, cell shape, membrane fluidity and chromosome organization (45-47). Treating cells with A22, a chemical that depolymerizes MreB, led to previously described aberrant cell morphologies (**Supplementary Movie 5**), yet our analysis showed that *tsr* diffusive behavior was unaltered compared to untreated cells (**Figure 2B**).

Taken together, our results suggest that reduction in membrane fluidity in a Δ*lpp*Δ*pgsA* strain affects mRNA diffusive behavior, but depolymerization of MreB does not.

### Translation of *tsr* mRNA occurs at hotspots determined by the physical location of the expressed gene

Next, we used the dynamic information gathered by live cell single-molecule tracking to visualize preferred mRNA localization sites depending on the molecules’ diffusive behavior. This allowed us to derive the distribution of active sites of *tsr* mRNA translation at the bacterial membrane, and thus gain understanding of the rules that govern their distribution at the subcellular level. To access the probability of finding an mRNA molecule at a certain location within the cell, we overlapped all the recorded trajectories, classified them according to the diffusion constants into “slow” (low-*D* and intermediate-*D* particles) and “fast” high-*D* particles, and projected them into the averaged normalized cell boundaries (see **Material and Methods** for details). Since we did not distinguish between the two bacterial cell poles, we “folded” cells in half to focus on the difference between polar and mid-cell localization densities. In contrast to a recent report that describes diffuse mRNA localization patterns in *B. subtilis* (21), we found that the slow and intermediate population of mRNA molecules displayed distinct localization patterns that we interpret as active translation sites, while fast subpopulations could be found anywhere in the bacterial cell with roughly the same frequency (**Figure 3**). To compare the distribution of translation sites with the location of the chromosome, we additionally imaged a fluorescent marker decorating the chromosome, HupA-mKate2, or stained chromosomes using Sytox Orange (48).

**Figure 3:**
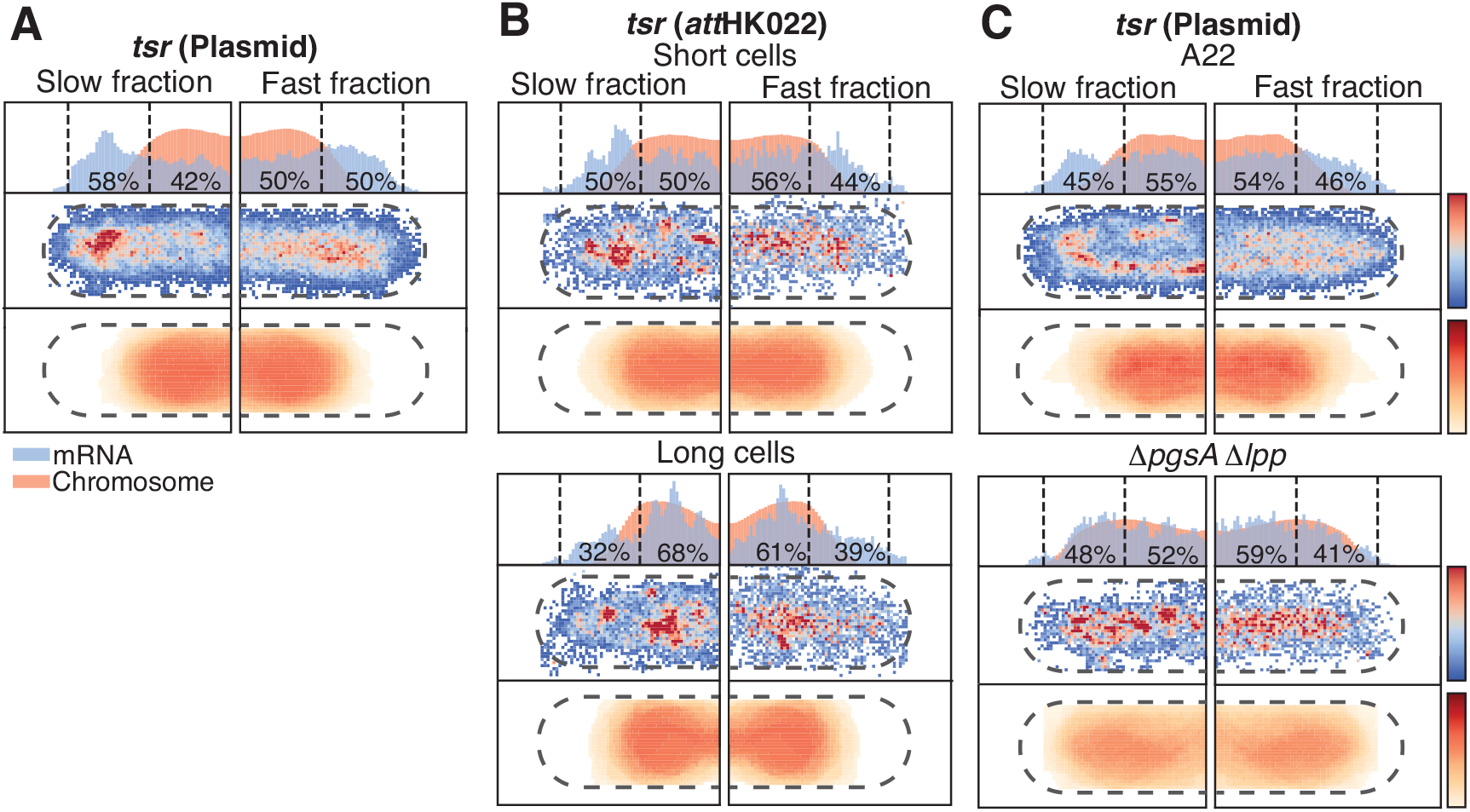
The slow and intermediate fraction of *tsr* mRNA localizes into hotspots whose location depends on the physical location of the transcribed gene and A22 treatment or removal of anionic phospholipids. a) Average localization maps of cell expressing *tsr* from pZA21 plasmid or b) from the *att*HK022 site on the bacterial chromosome near the terminus of replication. The middle panel shows the heat map of mRNA density for slow and intermediate (left) and fast (right) diffusive subpopulations from Figure 2. As there is no distinction between poles in our analysis, the densities are summed up into ‘folded’ cells. The direction of the cell halves is chosen for illustrative purposes. Bottom panel represents the averaged heat map of chromosome density based on HupA-mKate2 or Sytox Orange (Δ*lpp*Δ*pgsA* strain)) densities. The same cell “folding” method here results in symmetrical heat maps for left and right halves. The top panel shows 1-D histograms for the x axes on the two bottom panels. Numbers indicate the fraction of mRNA particles near the cell pole and cell center. b) Same as a) for *tsr* transcribed from the chromosome. The three top panels show the heat maps for cells smaller than the average cell size (short cells), the bottom three panels for larger than average cells (long cells). c) Average localization maps of cells expressing *tsr-24xMS2* from pZA21 plasmid treated with the MreB-depolymerizing agent A22 (top panel), and average localization maps of cells devoid of anionic phospholipids (Δ*lpp*Δ*pgsA)* expressing *tsr* from pZA21 plasmid (bottom panel). Numbers indicate the fraction of mRNA particles near the pole and cell center.

The slow and intermediate subpopulation of *tsr* mRNA expressed from a low copy-number plasmid (p15A *ori*) showed high densities at the edge of the nucleoid-occupied region, with only subtle differences between cells of different sizes (**Figure 3A, Supplementary Figure 4**). These observations follow the expectation of the subcellular distribution of passively segregated plasmids which were reported to localize at the edge of chromosome lobes (49). In contrast, the slow subpopulation of *tsr* expressed from a chromosomal single-copy insertion near the terminus of replication (*ter*) was distributed in punctate patterns overlapping with the chromosome-occupied region in small cells (cells smaller than the average population size), which presumably have a single copy of the chromosome. Cells larger than the population average, having replicated the chromosome, displayed a pronounced localization maximum near the cell center and decreased densities in the polar region, which is in good agreement with the localization of *ter* (50), and thus the *tsr* transcription source from *att*HK022 positioned at mid-cell (**Figure 3B**). The non-translating noAUG*tsr* mRNA construct was evenly distributed across the cell, while *tsr*Δ*1* displayed less defined localization patterns, and *tsr*Δ*2* appeared nearly evenly distributed within the cell (**Supplementary Figure 4**).

Cells treated with A22 displayed a drastic rearrangement of *tsr-24xms2* densities around the bacterial chromosome in a circular fashion (**Figure 3C**). A22 leads to loss of rod-shaped cellular morphology and consequently loss of cell polarity (45), which likely causes redistribution of plasmids along the edge of the chromosome and the observed change in densities. Δ*lpp*Δ*pgsA* cells also displayed aberrant chromosomes and cell shapes, and consequently altered *tsr-24xms2* distribution patterns. Nevertheless, in Δ*lpp*Δ*pgsA* cells, the distribution of slow *tsr-24xms2* mRNA subpopulations was distinctly different from A22 treated cells with a large fraction of slow-moving particles overlapping with the chromosome-occupied region (**Figure 3C**). SecYEG was shown to interact with and require APLs for proper function (42). Also, FtsY preferentially interacts with APLs in in-vitro studies (51). We thus conclude that the process of membrane targeting, and translation is severely disturbed in an APL mutant.

### Tsr expression affects the physical location of the *tsr* gene

To confirm that the observed spatiotemporal mRNA behavior is related to transertion, we visualized the genetic location of *tsr* expression using the *parS*-ParB system (52). According to the transertion model and previous observations (53), the source of gene expression displaces towards the cell membrane upon expression of a transmembrane protein. We labelled the plasmid expressing *tsr* and the chromosomal *tsr* insertion site with a *parS* sequence, and expressed a fluorescent mCherry-ParB at low levels driven by a constitutive promoter. We found that the *tsr-parS* plasmid showed some preference for cell poles but was overall evenly distributed within the cell (**Figure 4**). Upon induction of *tsr* expression using aTc, plasmid localization was profoundly altered, and the majority of mCherry-ParB-labelled plasmid was localized at the membrane into punctate patterns. Quantitative analysis confirmed that the plasmid was strongly displaced from the cell axis towards the cell periphery. Moreover, the plasmid showed increased densities near cell poles and thus the location of translation hotspots (**Figure 4 B**).

**Figure 4:**
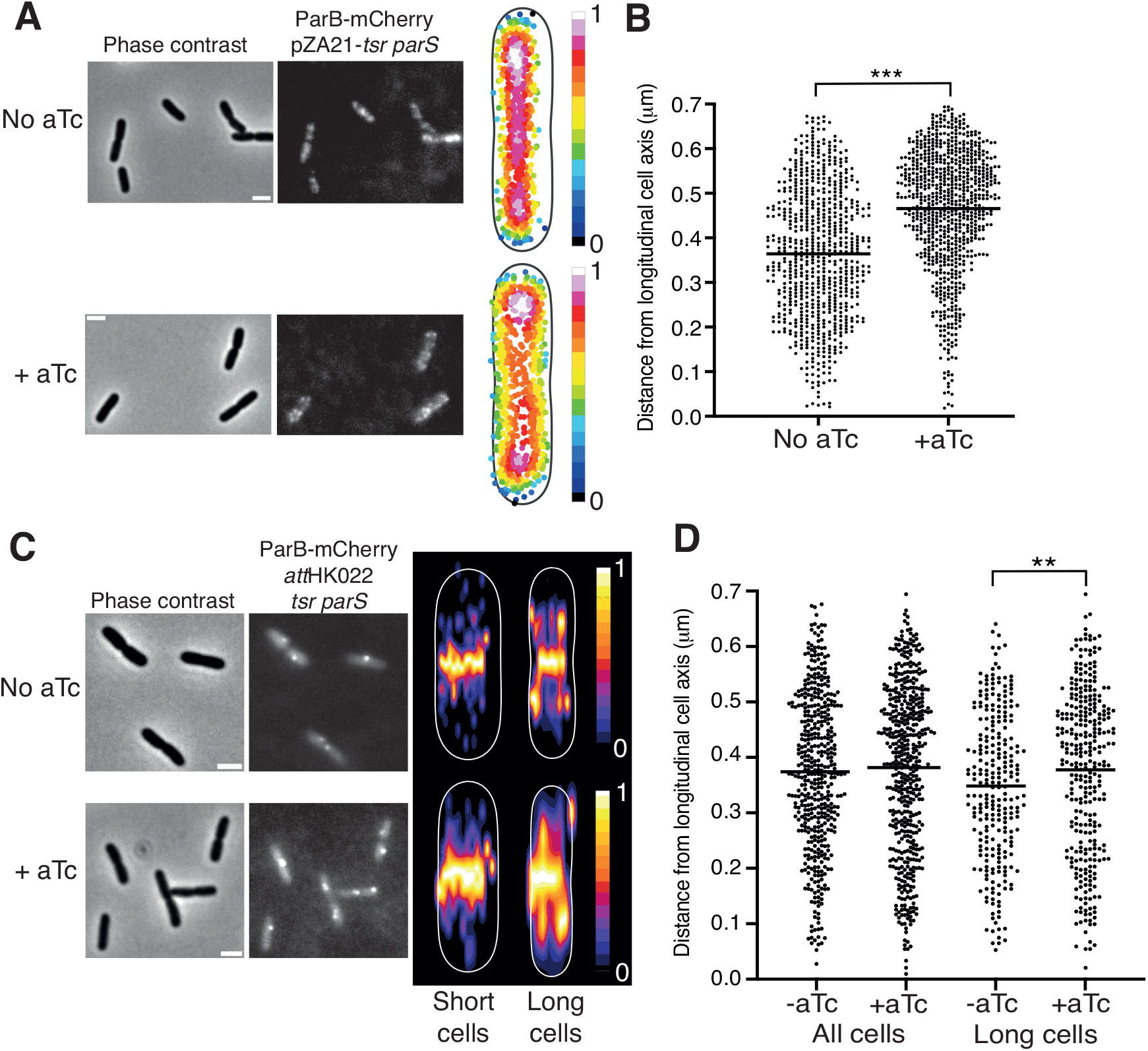
Localization of plasmid and *ter* site depends on *tsr* transcription. A) Localization of pZA21*-tsr parS* depends on *tsr* transcription. Top row: mCherry-ParB fluorescence shows diffuse to polar distribution of pZA21 within the cell with increased densities around cell poles in the absence of the inducer aTc in a normalized cell. Bottom row: In the presence of aTc, the mCherry-ParB signal is membrane-associated with increased densities near cell poles in a normalized cell. B) Quantification of pZA21*-tsr parS* displacement from the cell’s central axis connecting both cell poles upon induction with aTc. The mean distance of foci from the cell axis of uninduced cells is 0.35 *μ*m and 0.46 *μ*m for aTc-induced cells and statistically significant using a non-parametric Mann-Whitney test (p<0.0001). Sample sizes: 317 cells with 711 foci (no aTc), 276 cells with 814 foci (aTc). C) Localization of *att*HK02*2-tsr parS* depends on *tsr* transcription and follows the expectation of the location of *ter*. mCherry-ParB-labelled *att*HK022*-tsr parS* localizes to the cell center in small cells and additionally off-center in large cells. Cells were separated by the population average into small and large cells (2-3.7 *μ*m and 3.7-5.9 *μ*m, respectively). Upon induction with aTc, averaged cells show increased blurring around pronounced mid-cell densities. D) There is no difference in localization of *att*HK022*-tsr parS* between non-induced and induced cells. This difference becomes statistically significant if large cells (3.7-5.9 *μ*m) are binned and compared using a non-parametric Mann-Whitney test (p<0.008). Mean values are: all cells, 0.37 *μ*m (no aTc) and 0.38 *μ*m (+aTc). Large cells, 0.35 *μ*m (no aTc) and 0.38 *μ*m (+aTc). Sample sizes: no aTc, 551 cells, 447 foci. With aTc, 530 cells, 500 foci. Scale bars = 2*μ*m.

Fluorescent labelling of the chromosomal *att*HK022 site yielded fluorescent foci at mid-cell in small cells, and additional off-center densities in larger cells that had undergone full chromosome separation (**Figure 4C**), which follows our observation of hotspots in large cells and the expected position of *ter* in *E. coli* (50). Induction of *tsr* expression had only mild effects on displacement of *ter* and was more pronounced in large cells.

Overall, our results suggest that *tsr* mRNA membrane targeting and translation follows a basic proximity rule: the approximate location of the gene that encodes the expressed gene determines the distribution of *tsr* mRNA translation sites at the inner membrane, and *tsr* translation displaces the expressed gene towards the membrane. These observations are in good agreement with the process of transertion that describes physical coupling of transcription, translation and co-translational membrane protein insertion (53). Thus, we conclude that the translational hotspots we report here are a result of transertion and are formed in close proximity to the transcribed gene, which in turn is brought closer to the membrane during active protein translation.

## Discussion

A large fraction of the bacterial proteome is directly synthesized into the inner membrane of Gram negative bacteria, and this process shapes the overall spatial distribution of mRNA transcripts within the cell. Although highly dynamic in their nature, bacterial transcriptomes have mostly been studied in fixed cells. Here, we investigated and characterized the diffusive behavior of mRNA molecules encoding the Tsr chemoreceptor in order to uncover the rules governing transmembrane protein translation. In line with recent findings in the Gram-positive bacterium *Bacillus subtilis* (21), we found that the diffusion of individual mRNAs in *E. coli*, a Gram-negative organism, also displays characteristic subpopulations. The absence of the Sec-translocon-associated population in experiments with truncated *tsr* revealed a minimal polysome size required for strong membrane association. mRNA diffusion was slowed down by reduction of membrane fluidity in the APL-deficient strain. However, to our surprise, MreB depolymerization, assumed to also affect membrane fluidity, had no discernible effects on the distribution of diffusive states.

Having demonstrated that slow fractions of the diffusing mRNA most likely represented actively translated and membrane-bound transcripts, we uncovered that their translation occurred in hotspots, i.e. well-defined regions that are highly enriched in near-stationary mRNA particles, very similar to what has been reported in eukaryotes (20). Furthermore, the subcellular distribution of translation hotspots was non-random and dependent on the location of the underlying source of gene expression. Plasmid-encoded mRNA was preferentially translated between the nucleoid edge and the cell pole, while transcripts originating from the near-*ter* chromosomal copy of the *tsr* gene localized in the nucleoid-occupied part of the bacterial cell. In A22-treated cells and the APL mutant, the subcellular distribution of hotspots was strongly altered. As overall morphology is severely impacted in these cells, a combination of several effects including loss of cell shape and membrane defects affecting Sec function in the APL mutant may govern the observed redistribution of hotspots.

Directly visualizing the localization of the plasmid and the genomic locus of *tsr* expression confirmed their overlapping localization patterns, and localizations matched the distribution of hotspots at the membrane. More importantly, *tsr* expression led to reorganization and association of the plasmid with the membrane. Similarly, the genomic *tsr* locus was displaced towards the inner membrane, although this effect was less pronounced. Here, we hypothesize that the subcellular localization of the terminus region could be restricted by the workings of additional positioning or segregation mechanisms through FtsK (54) or MukBEF-MatP (55). Displacement of genomic loci and membrane-association of plasmids occurs as a function of transertion (53, 56), and based on these results, we suggest that hotspots occur as a product of transertion (53).

We propose a mechanism that determines membrane protein translation in proximity to the source of gene expression, which thus leads to the formation of hotspots. Co-localization of gene location, mRNA and transmembrane protein expression is likely determined by spatial constraints within the bacterial cell, where the nucleoid periphery or a plasmid is close to the inner membrane at any time. Therefore, the lifetime of 3D and 2D diffusion of *tsr* mRNA until binding to the membrane and Sec will be very short. Indeed, recent work investigating the kinetics of SRP shows an average search time of approximately 750 ms until binding to Sec occurs (57). This short search time will consequently restrict exploration of space of *tsr* mRNA until initiation of translation, leading to the observed overlap in gene location and translation. Following initiation of *tsr* translation, hotspot formation occurs as a function of mRNA length and formation of a transertion structure that includes localization of genetic loci or plasmid to the membrane. Furthermore, our findings suggest that multiple interactions between ribosomes and secretion systems are required for immobilization of *tsr* and the formation of defined hotspots. In turn, reduced ribosome binding increases mobility of shorter mRNA molecules, leading to less defined hotspots or no hotspots at all for very short transcripts.

Over two decades ago, it was proposed that long-lived transertion sites form hyperstructures, and our findings indicate the presence of membrane-associated hyperstructures that are driven by the process of translation (58). Our results connect the findings that transertion displaces genomic loci towards the membrane, leading to membrane heterogeneities driven by translation and co-translational membrane protein insertion (59). We hypothesize that translation hotspots, which have also been observed in eukaryotes (20), can foster protein-protein interactions and nucleation of multimeric protein clusters or large protein assemblies. For instance, operons encoding several transmembrane proteins forming a larger assembly are expected to be co-translated in close proximity, which would improve the efficiency of protein-protein interactions and complex assembly, as demonstrated to occur with cytosolic proteins in the bacterial cell (60).

## Acknowledgements & contribution

We thank Robert Hauschild and Ekaterina Papusheva and IST Austria Imaging Facility for support with TRIF microscopy and early-stage image analysis, and members of the Guet Group for helpful discussions. Ekaterina Krasnopeeva was funded by an EMBO fellowship, SS by an ERC Advanced Grant, and NN by the FWF (Austrian Science Fund) Elise Richter Program project number V 738.

TB and CCG planned study, and TB conducted experiments. TB, EK and SS analyzed data. NN provided materials for smFISH experiments. TB, EK and CCG wrote manuscript.

Raw mRNA tracking data is available at doi.org/10.5281/zenodo.7417870

## Material and Methods

### Media and Chemicals

All cloning and other standard molecular biology methods were done on strains grown in LB medium (Lennox, Sigma) and LB agar plates (1.5% agar, Sigma) supplemented with antibiotics as indicated. Defined minimal medium used for TIRF imaging consisted of 1xM9 salts, 0.05% casamino acid hydrolysate, 0.2% glucose, 1mM MgSO_4_ and 0.1mM CaCl_2_. All components were autoclaved separately, mixed and diluted to the indicated concentrations with sterile Millipore water, and sterile-filtered through a 0.22*μ*m filter before use. Antibiotics (Ampicillin, Chloramphenicol, Kanamycin, Spectinomycin, Rifampicin) were from Sigma, as were the inducers isopropyl-βD-thiogalactopyranosid (IPTG), anhydrous tetracycline (aTc) and the MreB inhibitor A22. SytoxOrange was from ThermoFisher Scientific. IPTG, aTc and A22 stocks were stored at −80°C. Small diluted aliquots were prepared and stored separately for imaging purposes and discarded after one freeze-thaw cycle.

We used the following chemicals for smFISH: nuclease-free deionized Formamide; 20xSSC, RNAse free; BSA, RNAse and DNAse free; 10xPBS, RNAse free; Diethylpyrocarbonate (DEPC)-treated water; all from Life Technologies/Ambion. Dextran sulfate sodium salt and tRNA from *E. coli* MRE600 (TRNAMRE-RO) was from Sigma Aldrich. Ribonucleoside-vanadyl complex (VRC) was from New England Biolabs. Formaldehyde was from Fisher Scientific.

### Strains, plasmids and enzymes used in this study

All strains used for imaging were derivatives of MG1655 unless indicated otherwise. All cloning was done in DH5a, its pir+ strain variant for cloning of CRIM plasmids (1), and in Frag1D. Frag1D is an *E. coli* Frag1B derivative carrying P_N25_-*tetR*, P_lacIq_-*lacI*-SpR in the lambda attachment site (2, 3), and a deletion of Δ*recA* introduced by lambda red recombineering. This strain was used to clone all pZA21 plasmids encoding MS2 stem loops with the purpose to repress transcription from P_LtetO1_, and to avoid loss of MS2 stem loop repeats by recombination.

Lambda red recombineering was done using pSIM6, pKD3, pKD4 and pKD13 PCR plasmids following established protocols (4). General P1 transduction was done using a previously described protocol (5). Single-copy insertion of plasmids with R6K origins of replication (CRIM plasmids) was done as described in (1). MG1655 and DH5α (DH5α pir+ or Frag1D) were grown at 37°C, strains harboring the plasmids pSIM6, pCP20 (6) or pAH69 (1) were grown at 30°.

Liquid cultures were grown at 250 rpm and 37°C or 30°C in 14ml plastic culture tubes (Falcon), and started by diluting overnight cultures 1:1000, which were yielded from single colony streaks. Strains were stored at −80°C as glycerol stocks in 15% glycerol.

All enzymes and DNA preparation kits used were from New England Biolabs (NEB), and all PCR products were amplified using *Phusion* polymerase. Restriction enzymes were HiFi versions where available, and DNA ligation was done with T4 DNA ligase at room temperature. Single-cut plasmid DNA was dephosphorylated using rSAP and gel-purified.

### Strain construction

All microscopy experiments were done using *E. coli* TB330 and its derivatives. This strain was constructed by sequentially introducing gene deletions or insertions into MG1655 (see list of strains used in this study **Table 2**). The first step was to introduce the P_N25_-*tetR*, P_lacIq_-*lacI*-SpR cassette from Frag1B (3) by P1 transduction into MG1655, followed by a P1 transduction introducing a PCR-verified *acrB*::*kan* deletion sourced from the KEIO collection (7) and curing the respective kanamycin resistance cassette using pCP20. As a last step, *recA* was deleted using lambda red recombineering followed by curing the resistance marker. Deletion of *acrB* was shown to ease induction of gene expression by P_LtetO1_using aTc by reducing aTc uptake heterogeneity (3), and deleting *recA* ensured stability of *24xms2* stem loop structures. We found that after one round of overnight culture of pZA21-*tsr-24xms2*, a large fraction of MS2 stem loops were lost, and that deleting *recA* abrogated or at least strongly minimized loss. To image the behavior of *tsr-24xms2* in a strain devoid of cardiolipin and phosphatidylglycerol, we further modified BKT25 (8), a strain deleted for *pgsA* and *lpp* with the P_N25_-*tetR*, P_lacIq_-*lacI*-SpR cassette and a *recA* gene deletion using P1 transduction. A HupA-mKate2 fusion at the endogenous *hupA* locus was constructed using lambda red recombineering and pNDL194 as PCR template (a gift from Nathan D. Lord, Harvard University, Boston). HupA was previously shown to be an excellent fusion partner to construct a constitutively expressed fluorescent chromosome label (9). HupA-mKate2 clones were verified by imaging, and a clone displaying a labelled chromosome was selected and used for all further experiments.

### Plasmid construction

To construct pZA21 variants, we amplified the *24xms2* cassette from pCR4-24xMS2SL-stable (Addgene #31865 (10)) such that the 5’ BamHI site of the *24xms2* cassette was removed and additional restriction sites were added at the 3’ end. The PCR product was cloned into pZA21 using KpnI and XbaI.

Cloning of all *tsr*–encoding plasmid variants was done by amplifying the *tsr* ORF or fragments from MG1655 genomic DNA. *tsr*Δ2 was ordered as gBlock from Integral DNA Technologies (IDT), Belgium. After digesting pZA21-*24xms2* and PCR fragments using KpnI, fragments were ligated and electroporated into electrocompetent Frag1D cells. Correct directionality of the inserts was verified by Sanger sequencing. A *tsr*-PAmCherry2 fusion was cloned by separately amplifying *tsr* and PAmCherry2 (Addgene #31932 from pPAmCherry2-C1) adding a GSGNKGQG polylinker between Tsr and PAmCherry2 and fusing both PCR products by fusion PCR followed by cloning into pZA21-*24xms2*. pZA21-*6xms2* was synthesized by Epoch Life Science, USA, and *tsr* was cloned using NheI and BsiWI. A *parS* site was added after opening pZA21-*tsr-24xms2* with HindIII followed by Gibson assembly with a *parS* gBlock (IDT). The sequence of the *parS* site is listed in **Table 4**.

To construct pZS*12-MS2(delFG N55K)-msfGFP (pZS*12-MS2-GFP hereafter), we amplified a MS2 delFG variant (3) lacking the delFG multimerization domain, and additionally introduced the high-affinity N55K mutation (11) by site-directed mutagenesis and fusion PCR. The corresponding PCR product was subsequently fused to msfGFP using fusion PCR, cloned into pZS*12-GFP replacing the GFP stuffer sequence, and sequence-verified by sequencing.

pAH68frt-cat-P_LtetO1_-*tsr-24xms2 parS* was constructed by digesting pAH68frt-cat (12) and pZA21*-tsr-24xms2* with SacI and BamHI and ligating the resulting pZA21*-tsr-24xms2* fragment lacking the p15A origin of replication with the pAH68frt-cat fragment. This plasmid was used to build TB378. To construct TB385, the pAH68frt-cat-P_LtetO1_-*tsr-24xms2* CRIM plasmid was digested with HindIII and combined with the *parS* sequence by ligation. After verification by sequencing, the plasmid was inserted into TB330 using pAH69, and the single copy state of the insertion was verified by colony PCR following protocols described in (1).

A fluorescent mCherry-ParB fusion was cloned by separately amplifying a ParB truncated for the first 30 amino acids (13) from a phage P1vir (Genbank AF234172) lysate and mCherry from (14). Both PCR products where combined using fusion PCR and a GSGNKGQG polylinker was added. The resulting PCR product was cloned into pZA32 with KpnI-HindIII. In the next step, the weak constitutive promoter J23114 (Anderson collection, iGem) was cloned into pZA32-mCherry-ParB replacing P_LlacO1_ between the XhoI and KpnI sites after annealing two phosphorylated complementary oligos encoding the promoter and containing matching sticky ends, resulting in plasmid pZA3-Pweak-mCherry-ParB. To image pZA21*-tsr-24xms2 parS*, we amplified JW23114-mCherry-ParB from pZA3-Pweak-mCherry-ParB and Gibson-assembled the fragment with a PCR product of pZS*12-MS2-GFP, placing JW23114-mCherry-ParB after MS2-GFP and the T1 terminator, resulting in pZS*12-MS2-GFP-Pweak-mCherry-ParB.

### Total internal reflection fluorescence microscopy

For TIRF microscopy, strains were grown to an OD_600nm_ of 0.2 in defined minimal medium which displayed low autofluorescence. 5 ng/ml aTc was added 15min prior to imaging of *tsr* mRNA particles and into the agar pad, and 5 *μ*M IPTG were supplied continuously during pre-culturing strains and during imaging to ensure low-level MS2-GFP expression.

25 *μ*g/ml Rifampicin and 10 ug/ml of the MreB inhibitor A22 were added to the indicated experiments 5 minutes before batch culture samples were prepared for imaging on agarose pads, to ensure complete hold of transcription and full depolymerization of MreB. Both Rifampicin and A22 were added into the agarose pad.

Staining of chromosomes in TB334 (Δ*lpp*Δ*pgsA*) using SytoxOrange was done as described elsewhere (15). We modified the protocol and used 50 nM instead of the reported 500 nM concentration. Furthermore, we applied the stain to the batch culture for 30 mins, washed away excess stain by pelleting cells and discarding the supernatant twice, followed by incubating the culture for another 1 h to allow dye dilution by growth and DNA replication.

Agarose pads were prepared by melting 2.5% agarose in minimal defined medium and by casting a pad between two glass slides spaced by two cover slips. 2×2 mm squares were cut out and placed into double-sided sticky 9×9 mm frame seals (Bio-Rad SLF0201) on a glass slide. 1 ml of bacterial culture at OD 0.3 were spun down, and the pellet was resuspended in 200 *μ*l of fresh low-autofluorescence medium. 1 *μ*l of this bacterial suspension was then spotted onto the agar pads and allowed to dry at room temperature for a few minutes before covering them with a high-precision and clean room grade cover slip (Schott Nexterion glass D, thickness 0.170+/- 0.005 mm).

Imaging was done at 30°C with a temperature-controlled Olympus IX81 total internal reflection fluorescence microscope equipped with a ZDC autofocus system, a water-cooled Hamamatsu Image EM C9100-13 camera, a 100×1.49NA objective lens and an additional 2x magnification tubular lens, resulting in a pixel size of 80 nm. The angle setting of the 488 nm diode laser was adjusted to a penetration depth of 80 nm to image MS2-sfGFP. Chromosomes labelled with HupA-mKate2 or SytoxOrange were imaged with the 561 nm laser and 350 nm penetration depth, and a single image was acquired before and after imaging MS2-GFP. Tsr-PAmCherry was photoactivated with short low-intensity pulses of the 405 nm laser.

The frame rate for membrane-associated *tsr-24xms2* mRNA and its shortened versions was 250 ms/frame, with an exposure time of 100 ms at 0.4mW 488 nm laser intensity, low camera gain settings and a readout speed of 2.75 MHz. *cfp-24xms2* and noAUG*tsr-24xms2* were imaged using 120 ms/frame and 60 ms/frame, respectively, a camera readout speed of 11 MHz and an oblique laser angle (HILO TIRF) setting of 350 nm. Individual XY positions were imaged up to 2000 frames, and up to 15 XY positions per agar pad were imaged in one imaging session.

### Image analysis and spot tracking

Kymographs were created using the ImageJ plugin “MultiKymograph” by drawing a segmented line of thickness 3 through the longitudinal axis of cells. Before kymographs were created, a mean filter of pixel size 1 was applied to images.

Single particle tracking was done using the ImageJ/FIJI plugin TrackMate (16) on manually outlined individual cell ROIs. The LoG or DOG detector was set to an “estimated blob diameter” of 0.45*μ*m with a threshold of 0.01, including a median filter and sub-pixel localization enabled. The initial thresholding of spots was chosen based on a quality criterion to minimize the number of false-positive spots and the result was visually inspected. The simple LAP tracker was chosen to connect spots into tracks without track splitting or merging allowed, with a linking max. distance and gap-closing distance of 0.5 *μ*m and no frame gaps allowed. In a final step, all tracks below 3 spots in track were discarded before exporting the tracks into a custom-written Python script.

The following mRNA variants were imaged only once: *24xms2, tsr-6xms2, tsr*Δ1-*24xms2*. All other constructs were imaged at least twice. HupA-mKate2 or SytoxOrange was imaged in separate experiments once for pZA21-*tsr-24xms2*, attHK022-*tsr-*24xms2, A22 pZA21-*tsr-24xms2* and TB334 (Δ*pgsA*Δ*lpp*) using SytoxOrange. Chromosome outlines were traced manually, saved as ROIs, and overlays were generated in Python.

### Diffusion coefficient calculations and distribution fitting

The xy positions of each particle for each frame were exported to .csv files and analyzed with custom written Python scripts using pandas and scipy (scipy.optimize and scipy.stats) packages. The mean square displacement for a single particle was calculated as:

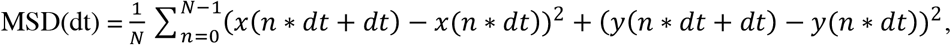

where N is the number of frames in a trajectory. The apparent diffusion coefficients were taken as the MSD(dt)/(4*dt) for dt = (1 time interval between frames) and plotted as histograms for all the trajectories recorded for a given condition.

The resulting distributions were fitted with the sums of Rayleigh distributions (from 1 to 4 terms) using scipy.optimize.curve_fit function. The Bayesian information criterion (BIC) was calculated as

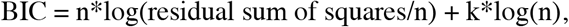

where n is a number of data points and k is a number of free parameters of the fitted model. A cutoff of 0.0005 um^2^/s was chosen to limit the near-zero fitting artefacts.

### Heat maps calculation

All the recorded cells from a given condition and their corresponding mRNA trajectories were re-oriented horizontally using coordinates transformation:

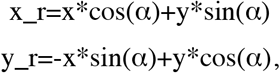

where x_r and y_r are rotated coordinates, x and y are original coordinates, and α is the angle of cell’s long axis rotation. The cell sizes were then normalized independently in x and y directions, and all the cells plus their trajectories were overlapped with the cell center placed at (0,0). The cells were then “folded” in half by taking absolute values of x.

Chromosomal heat maps were calculated in a similar way by overlapping all the chromosome ROI coordinates exported from Fiji/ImageJ.

### In-situ smFISH labelling of *tsr24xms2* mRNA

We closely followed a previously published protocol (17) to use single-molecule fluorescence in-situ hybridization (smFISH) to verify membrane localization of *tsr-24xms2* in the absence of MS2-GFP fusion protein, including probe design, cell preparation and probe concentration. We designed two probes that targeted the *24xms2* loops such that each probe would bind 12 times. Probes were ordered with Alexa 488 modification at 3’ from IDT. The sequences were

MS2 probe 1: TTCTTGGCAATAAGTACCGT

MS2 probe 2: ATGAACCCTGGAATACTGGA

We used 40% Formamide throughout. We resuspended cells in PBS before imaging, and imaging was done on agarose pads as described in the TIRF imaging section using a previously described Olympus IX81 microscope (18).

### Imaging the localization of pZA21 and att*HK022* using *parS*-ParB

To image the localization of pZA21 plasmid, we co-transformed pZA21*-tsr-24xms2 parS* and pZS*12-MS2-GFP-Pweak-mCherry-ParB into TB330. *tsr-24xms2* expression was induced using 10 ng/ml aTc, twice the concentration used for TIRF imaging, to guarantee homogenous induction across cells. pAH68frt-cat-P_LtetO1_-*tsr-24xms2 parS* inserted into att*HK022* was imaged by transforming TB385 with pZA3-Pweak-mCherry-ParB, and *tsr-24xms2* expression was induced with 20 ng/ml aTc, again to ensure homogeneity in induction. We grew, induced and placed cells on agarose pads as described for TIRF imaging. Imaging was done on an inverted Olympus IX83 that was identical to a previously described setup (18), but lacking a projector and having a Photometrix Prime 95B CMOS camera.

We analyzed images using the ImageJ plugin MicrobeJ (19) with a minimum cell size set to 1.8 *μ*m and maximal cell size to 6 *μ*m to capture cells with fully segregated ter, and a maximum cell width of 1.2 *μ*m. Parameters for foci detection in TB330 pZA21-*tsr-24xms2 parS* with or without induction of aTc was slightly more relaxed than for TB385 and fluorescently labelled ter. All lists of cells were manually corrected to exclude cells that contained abnormally high levels of fluorescence. To quantify pZA21-*tsr-24xms2 parS* localization, we dilated the segmentation mask by 250 nm to capture all spots at the cell periphery.

**Supplementary Table 1:** Predicted molecular weights of mRNA and proteins.

|  |  |
| --- | --- |
| *24xms2* | 500 kDa |
| MS2-GFP | 40 kDa (20, 21) |
| 24xMS2 decorated with 48xMs2-GFP | 1.8 MDa |
| *tsr* full length mRNA | 535 kDa |
| *tsr* Δ1 | 250 kDa |
| *tsr* Δ2 | 32.5 kDa |
| Bacterial ribosome | 2.7 MDa (22) |

**Supplementary Table 2:** Spot statistics.

| Condition | Number of cells | Number of trajectories |
| --- | --- | --- |
| *tsr* | 124 | 3481 |
|  | 52 of which with HupA-mKate2 labelled chromosomes (TB374) |  |
| *tsr* (attHK022) | 138 | 1670 |
|  | 57 of which with HupA-mKate2 labelled chromosomes (TB378) |  |
| *tsr-6xMS2* | 26 | 250 |
| *tsr*+Rifampicin | 63 | 636 |
| *noAUG-tsr* | 83 | 670 |
| *24xms2* | 31 | 109 |
| *cfp* | 41 | 418 |
| *lacY* | 37 | 306 |
| *tsrΔ1* | 47 | 628 |
| *tsrΔ2* | 38 | 336 |
| *tsr*+A22 | 103 | 2524 |
|  | 49 of which with HupA-mKate2 labelled chromosomes |  |
| *tsr ΔpgsAΔlpp* | 103 | 949 |
|  | 41 of which with SytoxOrange labelled chromosomes |  |

**Supplementary Table 3:**
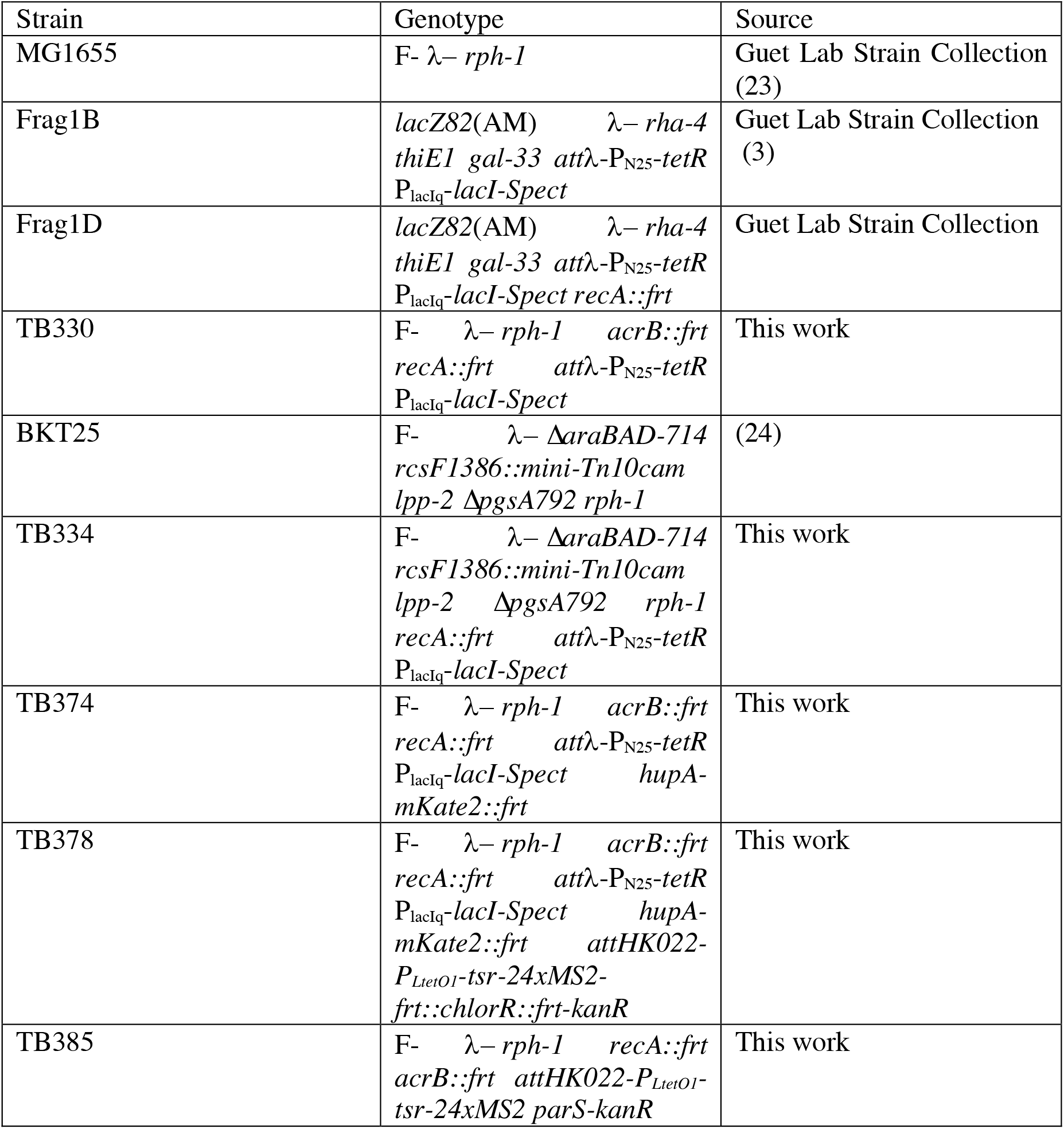
Relevant strains used in this study.

| Strain | Genotype | Source |
| --- | --- | --- |
| MG1655 | F- λ- *rph-1* | Guet Lab Strain Collection (23) |
| Frag1B | *lacZ82(AM)* λ- *rha-4 thiE1 gal-33 attλ-P<sub>N25</sub>-tetR P<sub>lacIq</sub>-lacI-Spect* | Guet Lab Strain Collection (3) |
| Frag1D | *lacZ82(AM)* λ- *rha-4 thiE1 gal-33 attλ-P<sub>N25</sub>-tetR P<sub>lacIq</sub>-lacI-Spect recA::frt* | Guet Lab Strain Collection |
| TB330 | F- λ- *rph-1 acrB::frt recA::frt attλ-P<sub>N25</sub>-tetR P<sub>lacIq</sub>-lacI-Spect* | This work |
| BKT25 | F- λ- Δ*araBAD-714 rcsF1386::mini-Tn10cam lpp-2 ΔpgsA792 rph-1* | (24) |
| TB334 | F- λ- Δ*araBAD-714 rcsF1386::mini-Tn10cam lpp-2 ΔpgsA792 rph-1 recA::frt attλ-P<sub>N25</sub>-tetR P<sub>lacIq</sub>-lacI-Spect* | This work |
| TB374 | F- λ- *rph-1 acrB::frt recA::frt attλ-P<sub>N25</sub>-tetR P<sub>lacIq</sub>-lacI-Spect hupA-mKate2::frt* | This work |
| TB378 | F- λ- *rph-1 acrB::frt recA::frt attλ-P<sub>N25</sub>-tetR P<sub>lacIq</sub>-lacI-Spect hupA-mKate2::frt attHK022-P<sub>LtetO1</sub>-tsr-24xMS2-frt::chlorR::frt-kanR* | This work |
| TB385 | F- λ- *rph-1 recA::frt acrB::frt attHK022-P<sub>LtetO1</sub>-tsr-24xMS2 parS-kanR* | This work |

**Supplementary Table 4:**
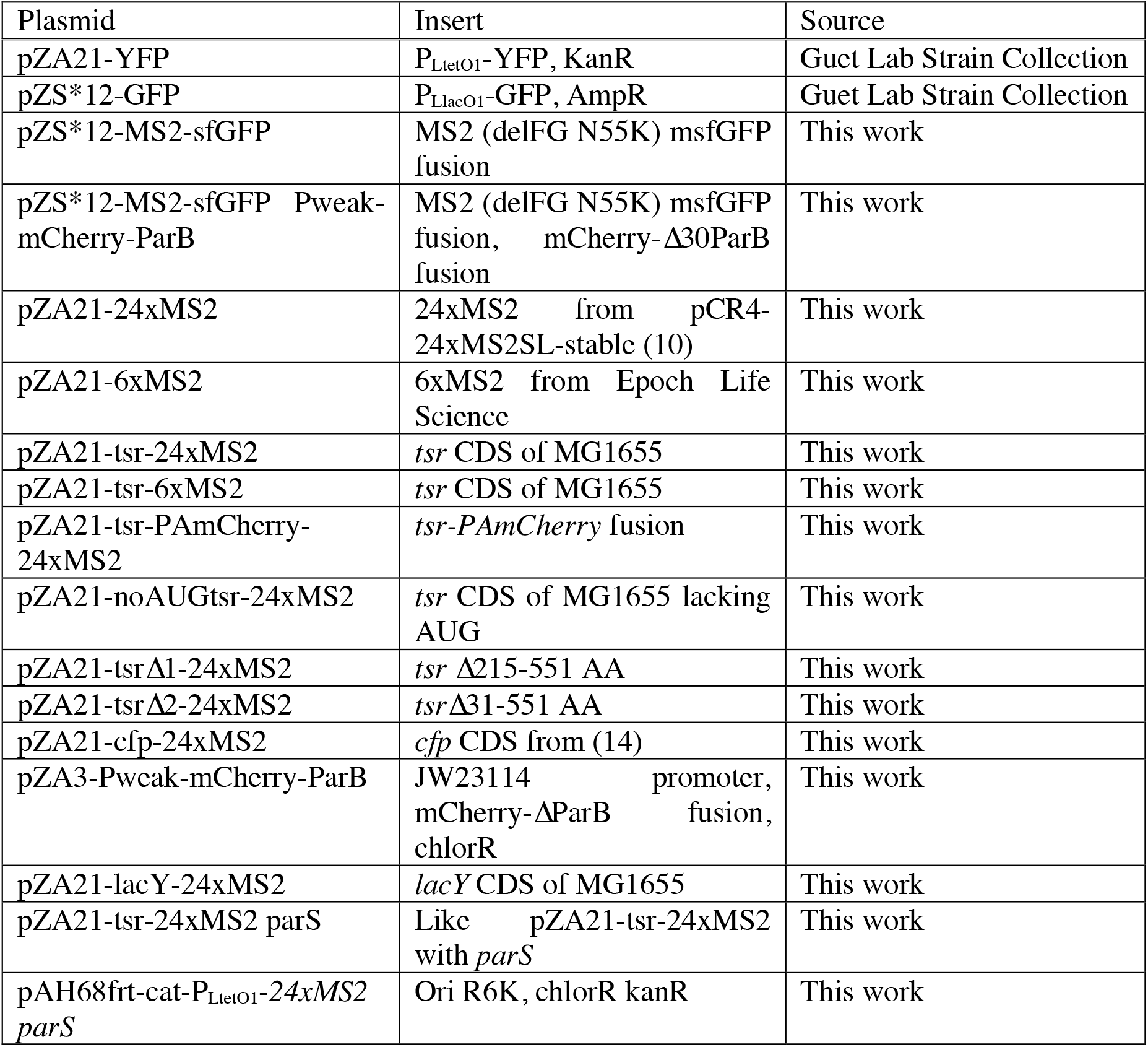
Relevant plasmids used in this study.

| Plasmid | Insert | Source |
| --- | --- | --- |
| pZA21-YFP | P<sub>LtetO1</sub>-YFP, KanR | Guet Lab Strain Collection |
| pZS*12-GFP | P<sub>LlacO1</sub>-GFP, AmpR | Guet Lab Strain Collection |
| pZS*12-MS2-sfGFP | MS2 (delFG N55K) msfGFP fusion | This work |
| pZS*12-MS2-sfGFP Pweak-mCherry-ParB | MS2 (delFG N55K) msfGFP fusion, mCherry-Δ30ParB fusion | This work |
| pZA21-24xMS2 | 24xMS2 from pCR4-24xMS2SL-stable (10) | This work |
| pZA21-6xMS2 | 6xMS2 from Epoch Life Science | This work |
| pZA21-tsr-24xMS2 | *tsr* CDS of MG1655 | This work |
| pZA21-tsr-6xMS2 | *tsr* CDS of MG1655 | This work |
| pZA21-tsr-PAmCherry-24xMS2 | *tsr*-PAmCherry fusion | This work |
| pZA21-noAUGtsr-24xMS2 | *tsr* CDS of MG1655 lacking AUG | This work |
| pZA21-tsrΔ1-24xMS2 | *tsr* Δ215-551 AA | This work |
| pZA21-tsrΔ2-24xMS2 | *tsr* Δ31-551 AA | This work |
| pZA21-cfp-24xMS2 | *cfp* CDS from (14) | This work |
| pZA3-Pweak-mCherry-ParB | JW23114 promoter, mCherry-ΔParB fusion, chlorR | This work |
| pZA21-lacY-24xMS2 | *lacY* CDS of MG1655 | This work |
| pZA21-tsr-24xMS2 parS | Like pZA21-tsr-24xMS2 with *parS* | This work |
| pAH68frt-cat-P<sub>LtetO1</sub>-24xMS2 *parS* | Ori R6K, chlorR kanR | This work |

**Supplementary Table 5:**
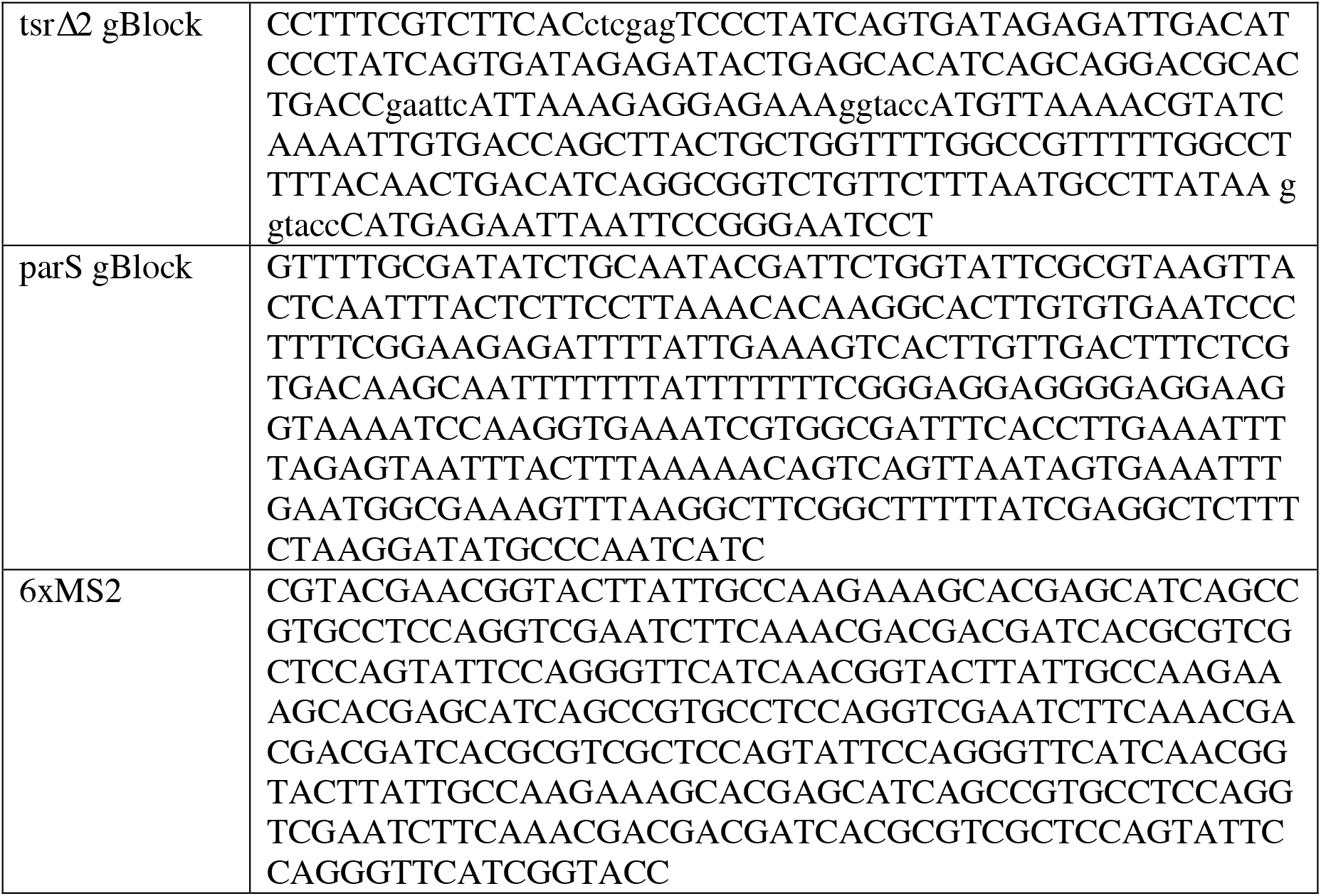
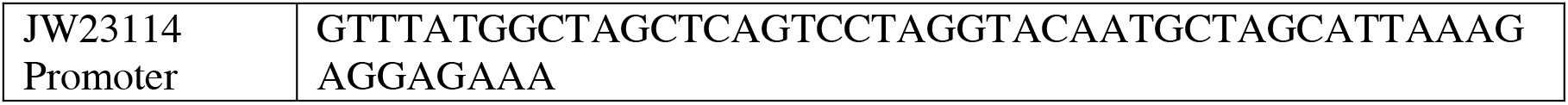
gBlocks and DNA sequences of 6xMS2 and JW23114.

**Supplementary Table 6:**
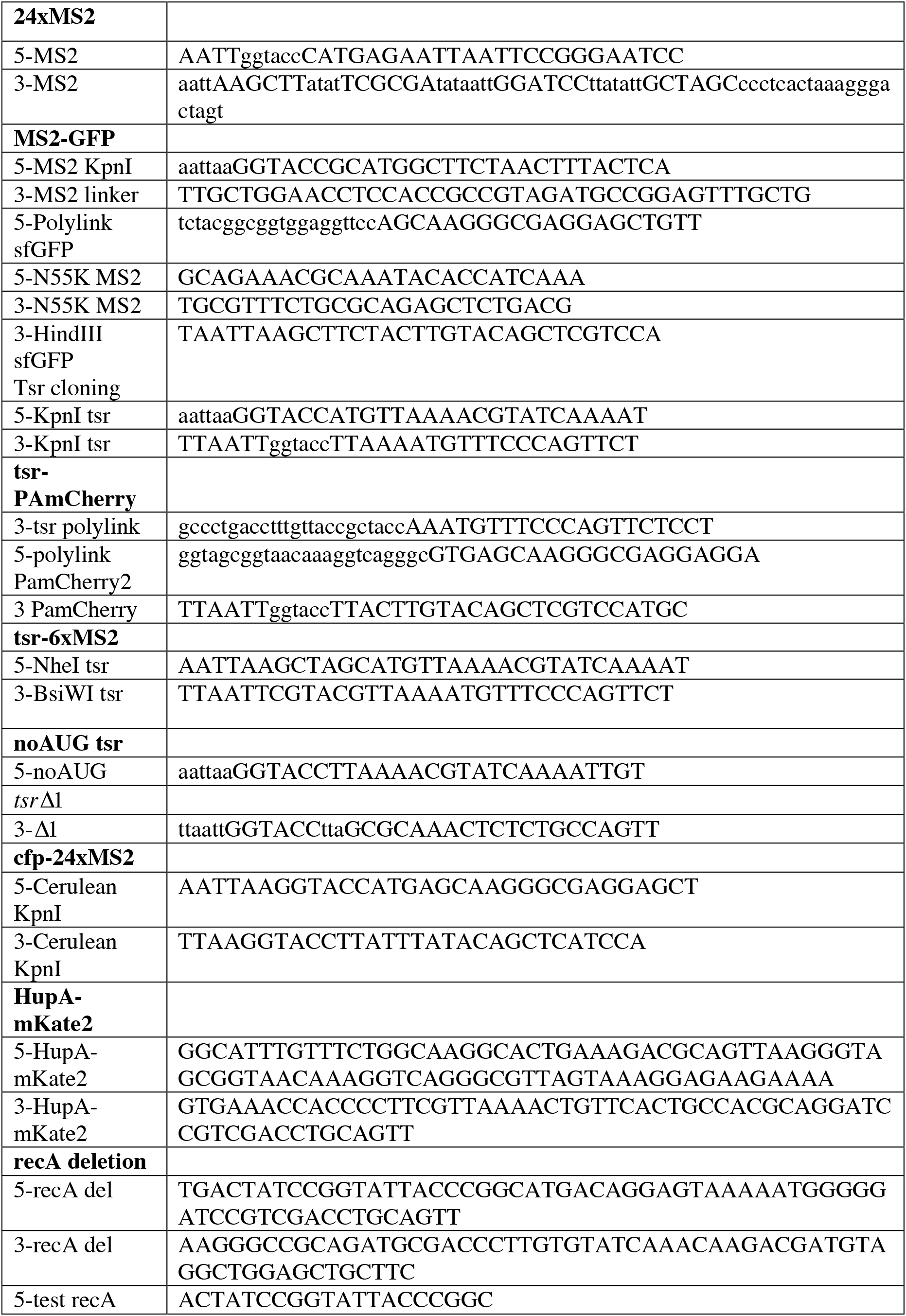
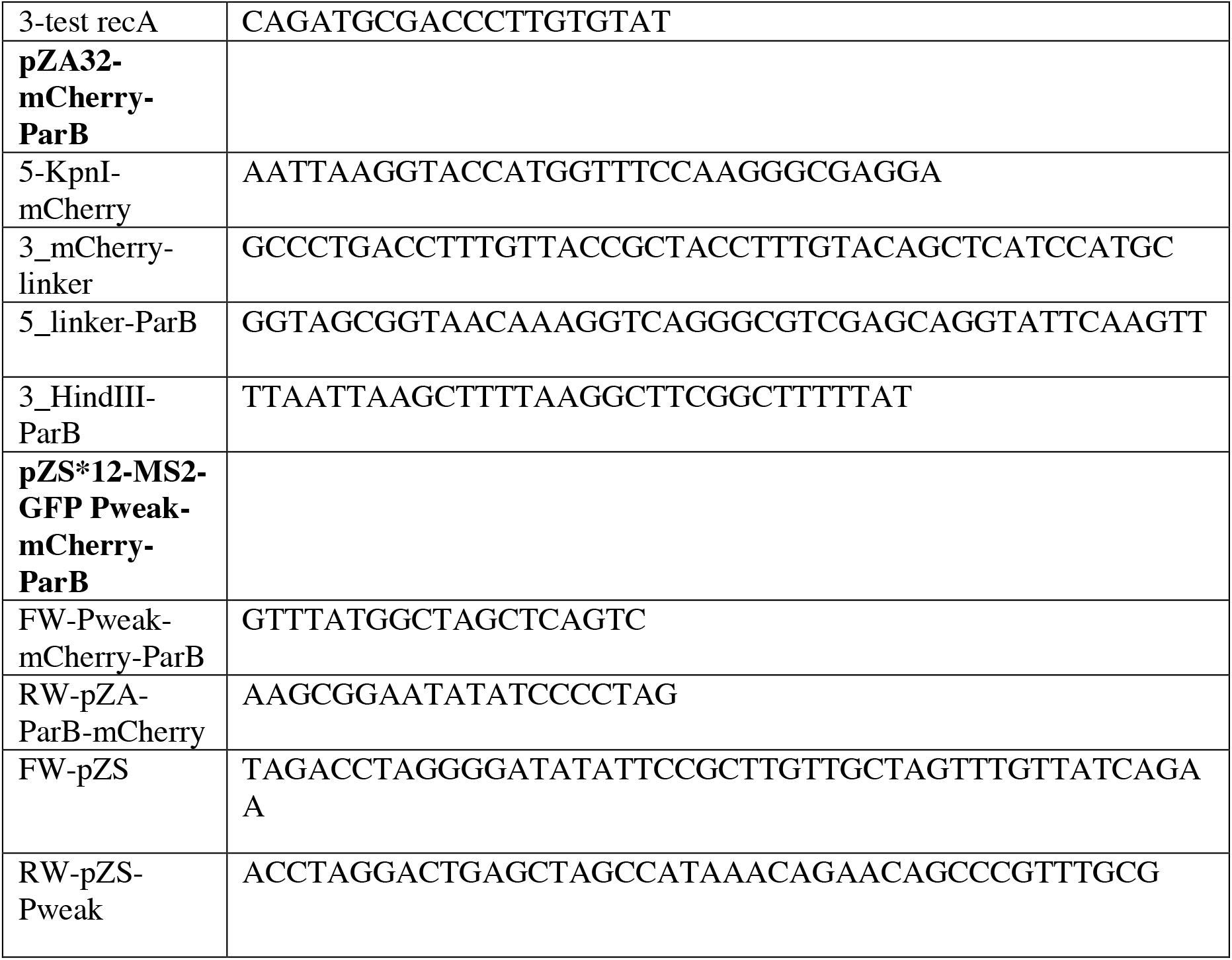
Oligonucleotides.

---

**Supplementary Figure 1.**
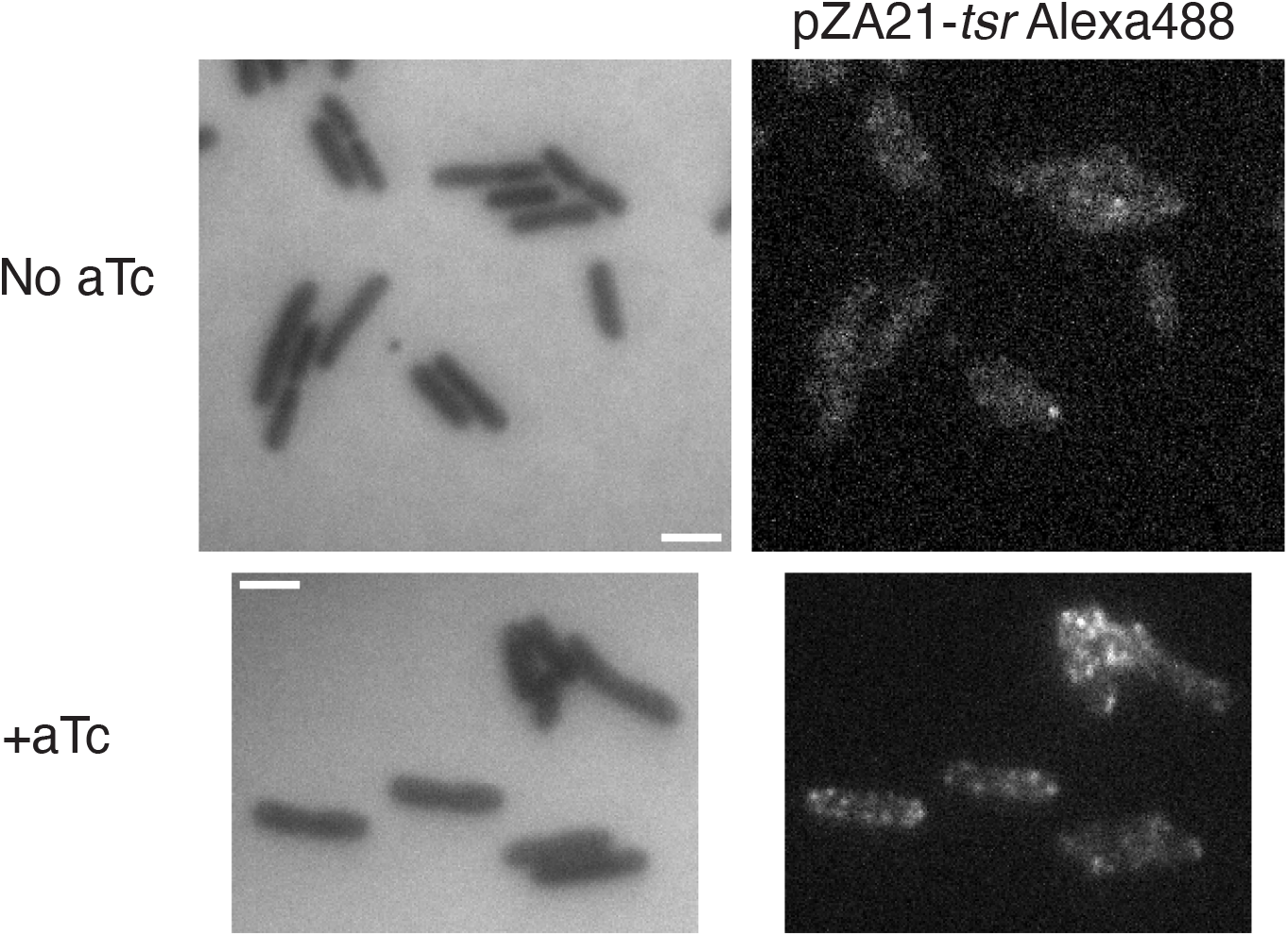
Images of cells stained using smFISH without (top) and with aTc (bottom). Left, cells imaged using reflected imaging mode. Right, Alexa 488 signal. mRNA localization appears in punctate patters along the membrane, suggesting that *tsr-24xms2* expressed from pZA21 is membrane-associated in the absence of MS2-GFP. Scale bar = 2*μ*m

**Supplementary Figure 2.**
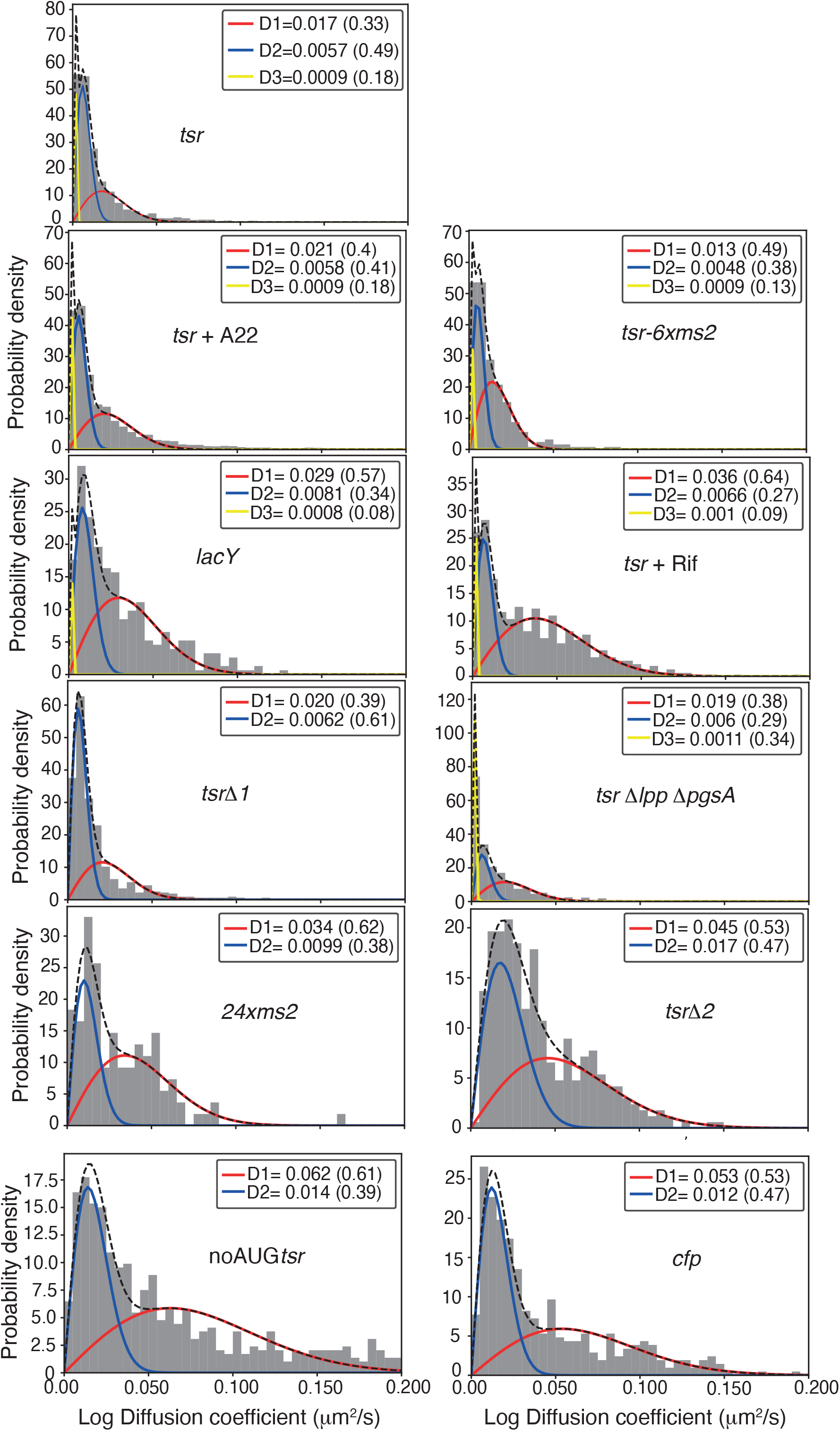
Single time step-derived diffusion coefficients distributions for all data shown in Figure 2 in the main text. mRNA type is highlighted within each histogram.

**Supplementary Figure 3.**
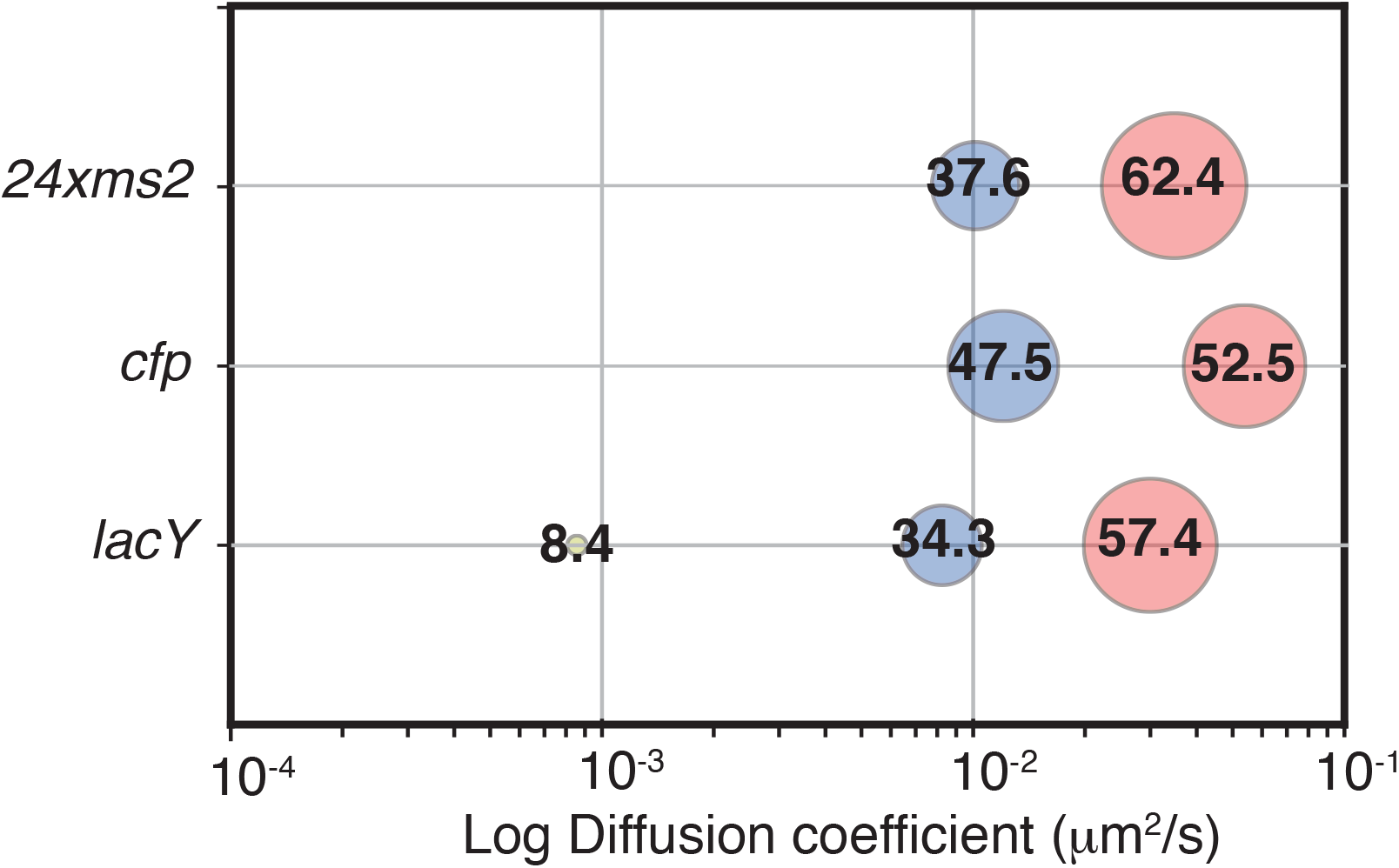
Bubble plot showing the summary of the fitting procedure for different mRNA species. Numbers on top of bubbles and bubble size indicate the weight of each subpopulation. X-axis: log diffusion coefficient, *μ*m^2^/s. *24xms2 –* mRNA molecule containing 24 MBS only and no coding mRNA. *cfp –* mRNA encoding cytoplasmic CFP protein, labelled with 24 MBS and expressed from a pZA21 plasmid. *cY –lacY* mRNA encoding membrane protein LacY, labelled with 24 MBS and expressed from a pZA21 plasmid.

**Supplementary Figure 4.**
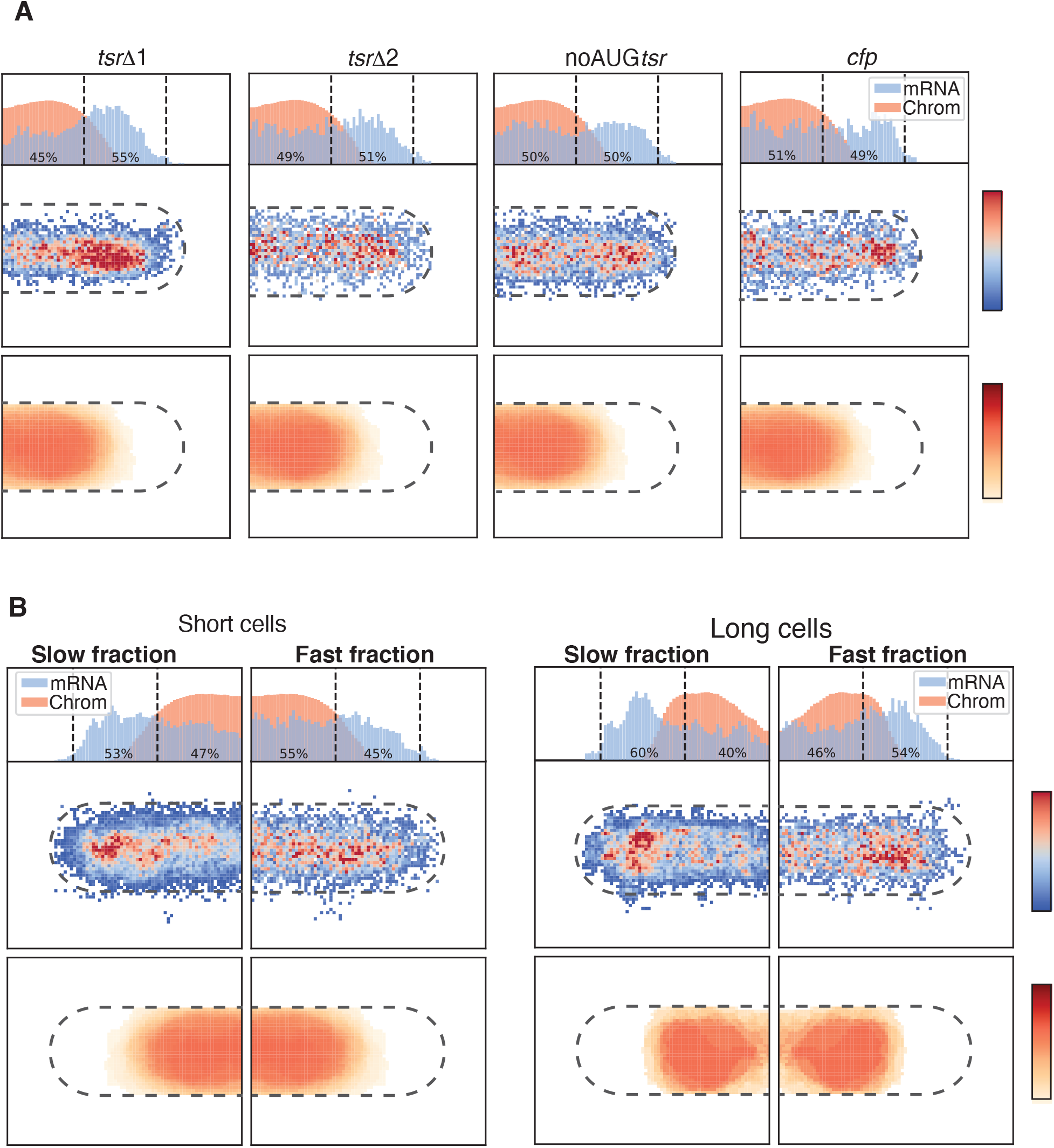
Average localization maps of cells expressing *tsr*Δ1*-24xms2, tsr*Δ2*-24xms2*, noAUG*tsr-24xms2* and *cfp-24xms2* from pZA21 plasmid. Intermediate and fast subpopulations were binned. b) Average localization maps of cells expressing *tsr-24xms2* from pZA21 sorted into short and long cells. Average chromosome localization derived from experiments with pZA21-*tsr-24xms2* cells was plotted into folded up cells and mirrored to the right. On top, probability density along the x-axis of the cell of mRNA particles and the chromosome is shown. Numbers indicate the fraction of mRNA particles near the cell pole and cell center. Slow subpopulations represent binned slow and intermediate populations.

## Supplementary Movies captions

**Supplementary Movie 1**

Cells expressing MS2-sfGFP only was image at 4Hz and compressed into 20fps. Scale bar = 2*μ*m

**Supplementary Movie 2**

Cells expressing *tsr-PAmCherry-24xms2*. MS2-sfGFP labelled mRNA was imaged at 4Hz and compressed into 20fps. Spot segmentation and tracking was done using TrackMate, tracks are uniformly color-coded. Tsr-PAmCherry was photoactivated using a short 405nm laser pulse at frame 201, 101 and 1 and imaged at 3Hz. Scale bar = 2*μ*m

**Supplementary Movie 3**

Cells expressing *tsr-24xms2*. MS2-sfGFP labelled mRNA was imaged at 4Hz and compressed into 20fps. Spot segmentation and tracking was done using TrackMate, tracks are uniformly color-coded. scale bar = 2*μ*m

**Supplementary Movie 4**

ΔlppΔpgsA cells with SytoxOrange stained chromosomes expressing *tsr-24xms2*. MS2-sfGFP labelled mRNA, imaged at 4Hz and compressed into 20fps. Spot segmentation and tracking was done using TrackMate, tracks are uniformly color-coded. A single 561nm image of Sytox Orange-stained chromosomes was taken after acquiring 488nm images. Scale bar = 2*μ*m

**Supplementary Movie 5**

Cells expressing *tsr-24xms2* and HupA-mKate2 and treated with A22. MS2-sfGFP labelled mRNA was imaged at 4Hz and compressed into 20fps. Spot segmentation and tracking was done using TrackMate, tracks are uniformly color-coded. A single 561nm image of HupA-mKate2-labelled chromosomes was taken after acquiring 488nm images. Scale bar = 2*μ*m

## Literature

1. N. Hadizadeh Yazdi, C. C. Guet, R. C. Johnson, J. F. Marko, Variation of the folding and dynamics of the Escherichia colichromosome with growth conditions. Molecular Microbiology 86, 1318–1333 (2012).

2. S. Bakshi, A. Siryaporn, M. Goulian, J. C. Weisshaar, Superresolution imaging of ribosomes and RNA polymerase in live Escherichia coli cells. Mol Microbiol 85, 21–38 (2012).

3. A. Sanamrad et al., Single-particle tracking reveals that free ribosomal subunits are not excluded from the Escherichia coli nucleoid. Proceedings of the National Academy of Sciences of the United States of America 111, 11413–11418 (2014).

4. X. Weng et al., Spatial organization of RNA polymerase and its relationship with transcription in Escherichia coli. Proc Natl Acad Sci U S A 116, 20115–20123 (2019).

5. D. Greenfield et al., Self-organization of the Escherichia coli chemotaxis network imaged with super-resolution light microscopy. PLoS biology 7, e1000137 (2009).

6. T. Bergmiller et al., Biased partitioning of the multidrug efflux pump AcrAB-TolC underlies long-lived phenotypic heterogeneity. Science 356, 311–315 (2017).

7. H. Strahl, J. Errington, Bacterial Membranes: Structure, Domains, and Function. Annu Rev Microbiol 71, 519–538 (2017).

8. M. Irastortza-Olaziregi, O. Amster-Choder, RNA localization in prokaryotes: Where, when, how, and why. Wiley Interdiscip Rev RNA 12, e1615 (2021).

9. M. Campos, C. Jacobs-Wagner, Cellular organization of the transfer of genetic information. Current opinion in microbiology 16, 171–176 (2013).

10. M. Castellana, S. Hsin-Jung Li, N. S. Wingreen, Spatial organization of bacterial transcription and translation. Proc Natl Acad Sci U S A 113, 9286–9291 (2016).

11. S. Park, M. Bujnowska, E. L. McLean, J. Fei, Quantitative Super-Resolution Imaging of Small RNAs in Bacterial Cells. Methods Mol Biol 1737, 199–212 (2018).

12. S. Kannaiah, J. Livny, O. Amster-Choder, Spatiotemporal Organization of the E. coli Transcriptome: Translation Independence and Engagement in Regulation. Mol Cell 76, 574–589 e577 (2019).

13. J. R. Moffitt, S. Pandey, A. N. Boettiger, S. Y. Wang, X. W. Zhuang, Spatial organization shapes the turnover of a bacterial transcriptome. Elife 5 (2016).

14. D. Akopian, K. Shen, X. Zhang, S. O. Shan, Signal recognition particle: an essential protein-targeting machine. Annu Rev Biochem 82, 693–721 (2013).

15. R. Steinberg, L. Knupffer, A. Origi, R. Asti, H. G. Koch, Co-translational protein targeting in bacteria. Fems Microbiology Letters 365 (2018).

16. S. Angelini, S. Deitermann, H. G. Koch, FtsY, the bacterial signal-recognition particle receptor, interacts functionally and physically with the SecYEG translocon. EMBO Rep 6, 476–481 (2005).

17. J. Frauenfeld et al., Cryo-EM structure of the ribosome-SecYE complex in the membrane environment. Nat Struct Mol Biol 18, 614–621 (2011).

18. M. Roggiani, M. Goulian, Chromosome-Membrane Interactions in Bacteria. Annu Rev Genet 49, 115–129 (2015).

19. K. Matsumoto, H. Hara, I. Fishov, E. Mileykovskaya, V. Norris, The membrane: transertion as an organizing principle in membrane heterogeneity. Frontiers in Microbiology 6 (2015).

20. Z. B. Katz et al., Mapping translation ‘hot-spots’ in live cells by tracking single molecules of mRNA and ribosomes. Elife 5 (2016).

21. L. Sattler, P. L. Graumann, Real-Time Messenger RNA Dynamics in Bacillus subtilis. Front Microbiol 12, 760857 (2021).

22. I. Golding, E. C. Cox, RNA dynamics in live Escherichia coli cells. Proceedings of the National Academy of Sciences of the United States of America 101, 11310–11315 (2004).

23. J. F. Gebert, B. Overhoff, M. D. Manson, W. Boos, The Tsr Chemosensory Transducer of Escherichia-Coli Assembles into the Cytoplasmic Membrane Via a Seca-Dependent Process. Journal of Biological Chemistry 263, 16652–16660 (1988).

24. E. Bertrand et al., Localization of ASH1 mRNA particles in living yeast. Molecular Cell 2, 437–445 (1998).

25. A. R. Buxbaum, G. Haimovich, R. H. Singer, In the right place at the right time: visualizing and understanding mRNA localization. Nature Reviews Molecular Cell Biology 10.1038/nrm3918 (2014).

26. C. C. Guet et al., Minimally invasive determination of mRNA concentration in single living bacteria. Nucleic Acids Research 36 (2008).

27. F. Lim, M. Spingola, D. S. Peabody, Altering the RNA binding specificity of a translational repressor. J Biol Chem 269, 9006–9010 (1994).

28. J.-D. Pédelacq, S. Cabantous, T. Tran, T. C. Terwilliger, G. S. Waldo, Engineering and characterization of a superfolder green fluorescent protein. Nature Biotechnology 24, 79–88 (2005).

29. R. Lutz, H. Bujard, Independent and tight regulation of transcriptional units in Escherichia coli via the LacR/O, the TetR/O and AraC/I1-I2 regulatory elements. Nucleic Acids Research 25, 1203–1210 (1997).

30. E. Tutucci et al., An improved MS2 system for accurate reporting of the mRNA life cycle. Nat Methods 15, 81–89 (2018).

31. A. Deana, J. G. Belasco, Lost in translation: the influence of ribosomes on bacterial mRNA decay. Genes Dev 19, 2526–2533 (2005).

32. D. W. Selinger, R. M. Saxena, K. J. Cheung, G. M. Church, C. Rosenow, Global RNA half-life analysis in Escherichia coli reveals positional patterns of transcript degradation. Genome Res 13, 216–223 (2003).

33. I. Andreeva, R. Belardinelli, M. V. Rodnina, Translation initiation in bacterial polysomes through ribosome loading on a standby site on a highly translated mRNA. Proc Natl Acad Sci U S A 115, 4411–4416 (2018).

34. M. Kumar, M. S. Mommer, V. Sourjik, Mobility of Cytoplasmic, Membrane, and DNA-Binding Proteins in Escherichia coli. Biophysj 98, 552–559 (2010).

35. P. Milon, C. Maracci, L. Filonava, C. O. Gualerzi, M. V. Rodnina, Real-time assembly landscape of bacterial 30S translation initiation complex. Nat Struct Mol Biol 19, 609–615 (2012).

36. K. K. Kim, H. Yokota, S. H. Kim, Four-helical-bundle structure of the cytoplasmic domain of a serine chemotaxis receptor. Nature 400, 787–792 (1999).

37. N. R. Voss, M. Gerstein, T. A. Steitz, P. B. Moore, The geometry of the ribosomal polypeptide exit tunnel. J Mol Biol 360, 893–906 (2006).

38. T. Yamamoto, S. Izumi, K. Gekko, Mass spectrometry of hydrogen/deuterium exchange in 70S ribosomal proteins from E. coli. FEBS Lett 580, 3638–3642 (2006).

39. M. Gohrbandt et al., Low membrane fluidity triggers lipid phase separation and protein segregation in living bacteria. EMBO J 41, e109800 (2022).

40. P. M. Oliver et al., Localization of anionic phospholipids in Escherichia coli cells. Journal of bacteriology 196, 3386–3398 (2014).

41. A. Nenninger et al., Independent mobility of proteins and lipids in the plasma membrane of Escherichia coli. Molecular Microbiology 92, 1142–1153 (2014).

42. S. Ryabichko et al., Cardiolipin is required in vivo for the stability of bacterial translocon and optimal membrane protein translocation and insertion. Sci Rep 10, 6296 (2020).

43. B. K. Tan et al., Discovery of a cardiolipin synthase utilizing phosphatidylethanolamine and phosphatidylglycerol as substrates. Proc Natl Acad Sci U S A 109, 16504–16509 (2012).

44. E. C. Garner et al., Coupled, Circumferential Motions of the Cell Wall Synthesis Machinery and MreB Filaments in B. subtilis. Science 333, 222–225 (2011).

45. T. S. Ursell et al., Rod-like bacterial shape is maintained by feedback between cell curvature and cytoskeletal localization. Proc Natl Acad Sci U S A 111, E1025–1034 (2014).

46. H. Strahl, F. Burmann, L. W. Hamoen, The actin homologue MreB organizes the bacterial cell membrane. Nature communications 5 (2014).

47. F. Oswald, A. Varadarajan, H. Lill, E. J. G. Peterman, Y. J. M. Bollen, MreB-Dependent Organization of the E-coli Cytoplasmic Membrane Controls Membrane Protein Diffusion. Biophysical journal 110, 1139–1149 (2016).

48. S. Bakshi et al., Nonperturbative imaging of nucleoid morphology in live bacterial cells during an antimicrobial peptide attack. Appl Environ Microbiol 80, 4977–4986 (2014).

49. Y. F. Li, S. Austin, The P1 plasmid is segregated to daughter cells by a ‘capture and ejection’ mechanism coordinated with Escherichia coli cell division. Molecular Microbiology 46, 63–74 (2002).

50. I. F. Lau et al., Spatial and temporal organization of replicating Escherichia coli chromosomes. Mol Microbiol 49, 731–743 (2003).

51. R. Parlitz et al., Escherichia coli Signal Recognition Particle Receptor FtsY Contains an Essential and Autonomous Membrane-binding Amphipathic Helix. The Journal of biological chemistry 282, 32176–32184 (2007).

52. T. B. K. Le, M. T. Laub, New approaches to understanding the spatial organization of bacterial genomes. Current Opinion in Microbiology 22, 15–21 (2014).

53. E. A. Libby, M. Roggiani, M. Goulian, Membrane protein expression triggers chromosomal locus repositioning in bacteria. Proceedings of the National Academy of Sciences of the United States of America 109, 7445–7450 (2012).

54. D. J. Sherratt, L. K. Arciszewska, E. Crozat, J. E. Graham, I. Grainge, The Escherichia coli DNA translocase FtsK. Biochem Soc Trans 38, 395–398 (2010).

55. J. Makela, D. J. Sherratt, Organization of the Escherichia coli Chromosome by a MukBEF Axial Core. Mol Cell 78, 250–260 e255 (2020).

56. A. S. Lynch, J. C. Wang, Anchoring of DNA to the bacterial cytoplasmic membrane through cotranscriptional synthesis of polypeptides encoding membrane proteins or proteins for export: a mechanism of plasmid hypernegative supercoiling in mutants deficient in DNA topoisomerase I. J Bacteriol 175, 1645–1655 (1993).

57. I. L. Volkov et al., Spatiotemporal kinetics of the SRP pathway in live E. coli cells. Proc Natl Acad Sci U S A 119, e2204038119 (2022).

58. V. Norris, M. S. Madsen, Autocatalytic gene expression occurs via transertion and membrane domain formation and underlies differentiation in bacteria: a model. J Mol Biol 253, 739–748 (1995).

59. D. Lopez, G. Koch, Exploring functional membrane microdomains in bacteria: an overview. Curr Opin Microbiol 36, 76–84 (2017).

60. Y. W. Shieh et al., Operon structure and cotranslational subunit association direct protein assembly in bacteria. Science 350, 678–680 (2015).

## Literature

1. A. Haldimann, B. Wanner, Conditional-replication, integration, excision, and retrieval plasmid-host systems for gene structure-function studies of bacteria. Journal of bacteriology 183, 6384–6393 (2001).

2. R. Lutz, H. Bujard, Independent and tight regulation of transcriptional units in Escherichia coli via the LacR/O, the TetR/O and AraC/I1-I2 regulatory elements. Nucleic Acids Research 25, 1203–1210 (1997).

3. T. T. Le et al., Real-time RNA profiling within a single bacterium. Proc Natl Acad Sci U S A 102, 9160–9164 (2005).

4. L. Thomason et al., Recombineering: genetic engineering in bacteria using homologous recombination. Curr Protoc Mol Biol Chapter 1, Unit 1.16 (2007).

5. E. S. Lennox, Transduction of linked genetic characters of the host by bacteriophage P1. Virology 1, 190–206 (1955).

6. K. A. Datsenko, B. L. Wanner, One-step inactivation of chromosomal genes in Escherichia coli K-12 using PCR products. Proc Natl Acad Sci U S A 97, 6640–6645 (2000).

7. T. Baba et al., Construction of Escherichia coli K-12 in-frame, single-gene knockout mutants: the Keio collection. Mol Syst Biol 2, 2006.0008 (2006).

8. B. K. Tan, M. Bogdanov, J. Zhao (2012) Discovery of a cardiolipin synthase utilizing phosphatidylethanolamine and phosphatidylglycerol as substrates. in Proceedings of the ….

9. F. Wu, E. Van Rijn, B. G. Van Schie, J. E. Keymer, C. Dekker, Multi-color imaging of the bacterial nucleoid and division proteins with blue, orange, and near-infrared fluorescent proteins. Front Microbiol 6, 607 (2015).

10. E. Bertrand et al., Localization of ASH1 mRNA particles in living yeast. Molecular Cell 2, 437–445 (1998).

11. F. Lim, M. Spingola, D. S. Peabody, Altering the RNA binding specificity of a translational repressor. J Biol Chem 269, 9006–9010 (1994).

12. T. Bergmiller et al., Biased partitioning of the multidrug efflux pump AcrAB-TolC underlies long-lived phenotypic heterogeneity. Science 356, 311–315 (2017).

13. H. J. Nielsen, J. R. Ottesen, B. Youngren, S. J. Austin, F. G. Hansen, The Escherichia coli chromosome is organized with the left and right chromosome arms in separate cell halves. Mol Microbiol 62, 331–338 (2006).

14. R. Cox, M. J. Dunlop, M. B. Elowitz, A synthetic three-color scaffold for monitoring genetic regulation and noise. Journal of Biological Engineering 4, 10 (2010).

15. S. Bakshi et al., Nonperturbative imaging of nucleoid morphology in live bacterial cells during an antimicrobial peptide attack. Appl Environ Microbiol 80, 4977–4986 (2014).

16. J. Y. Tinevez et al., TrackMate: An open and extensible platform for single-particle tracking. Methods 115, 80–90 (2017).

17. S. O. Skinner, L. A. S. u. lveda, H. Xu, I. Golding, Measuring mRNA copy number in individual Escherichia coli cells using single-molecule fluorescent in situ hybridization. Nature Protocols 8, 1100–1113 (2013).

18. R. Chait, J. Ruess, T. Bergmiller, G. Tkacik, C. C. Guet, Shaping bacterial population behavior through computer-interfaced control of individual cells. Nature communications 8 (2017).

19. A. Ducret, E. M. Quardokus, Y. V. Brun, MicrobeJ, a tool for high throughput bacterial cell detection and quantitative analysis. Nat Microbiol 1, 16077 (2016).

20. D. A. Kuzmanovic, I. Elashvili, C. Wick, C. O’Connell, S. Krueger, Bacteriophage MS2: molecular weight and spatial distribution of the protein and RNA components by small-angle neutron scattering and virus counting. Structure 11, 1339–1348 (2003).

21. W. Uckert, L. Pedersen, W. Günzburg, “Green Fluorescent Protein Retroviral Vector: Generation of High-Titer Producer Cells and Virus Supernatant”. (Humana Press), 10.1385/1-59259-086-1:275, pp. 275–285.

22. T. Yamamoto, S. Izumi, K. Gekko, Mass spectrometry of hydrogen/deuterium exchange in 70S ribosomal proteins from E. coli. FEBS Lett 580, 3638–3642 (2006).

23. F. R. Blattner et al., The complete genome sequence of Escherichia coli K-12. Science 277, 1453–1462 (1997).

24. B. K. Tan et al., Discovery of a cardiolipin synthase utilizing phosphatidylethanolamine and phosphatidylglycerol as substrates. Proc Natl Acad Sci U S A 109, 16504–16509 (2012).

