## Supplementary Figures for "Real-time dynamics of individual chemoreceptor mRNA molecules reveals translation hotspots at the inner membrane of *Escherichia coli*"

### Supplementary Figure 1

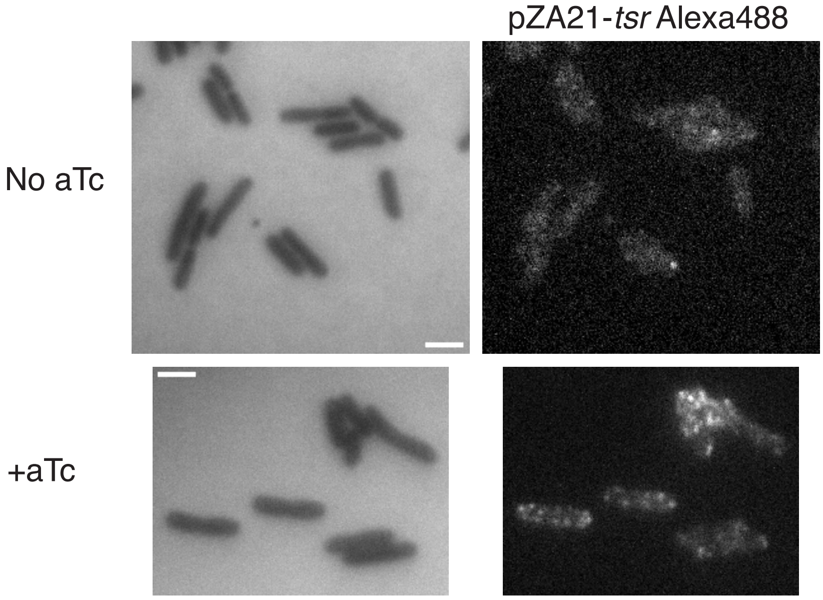


**Supplementary Figure 1**
Images of cells stained using smFISH without (top) and with aTc (bottom). Left, cells imaged using reflected imaging mode. Right, Alexa 488 signal. mRNA localization appears in punctate patters along the membrane, suggesting that *tsr-24xms2* expressed from pZA21 is membrane-associated in the absence of MS2-GFP. Scale bar = 2$\mu$m

### Supplementary Figure 2

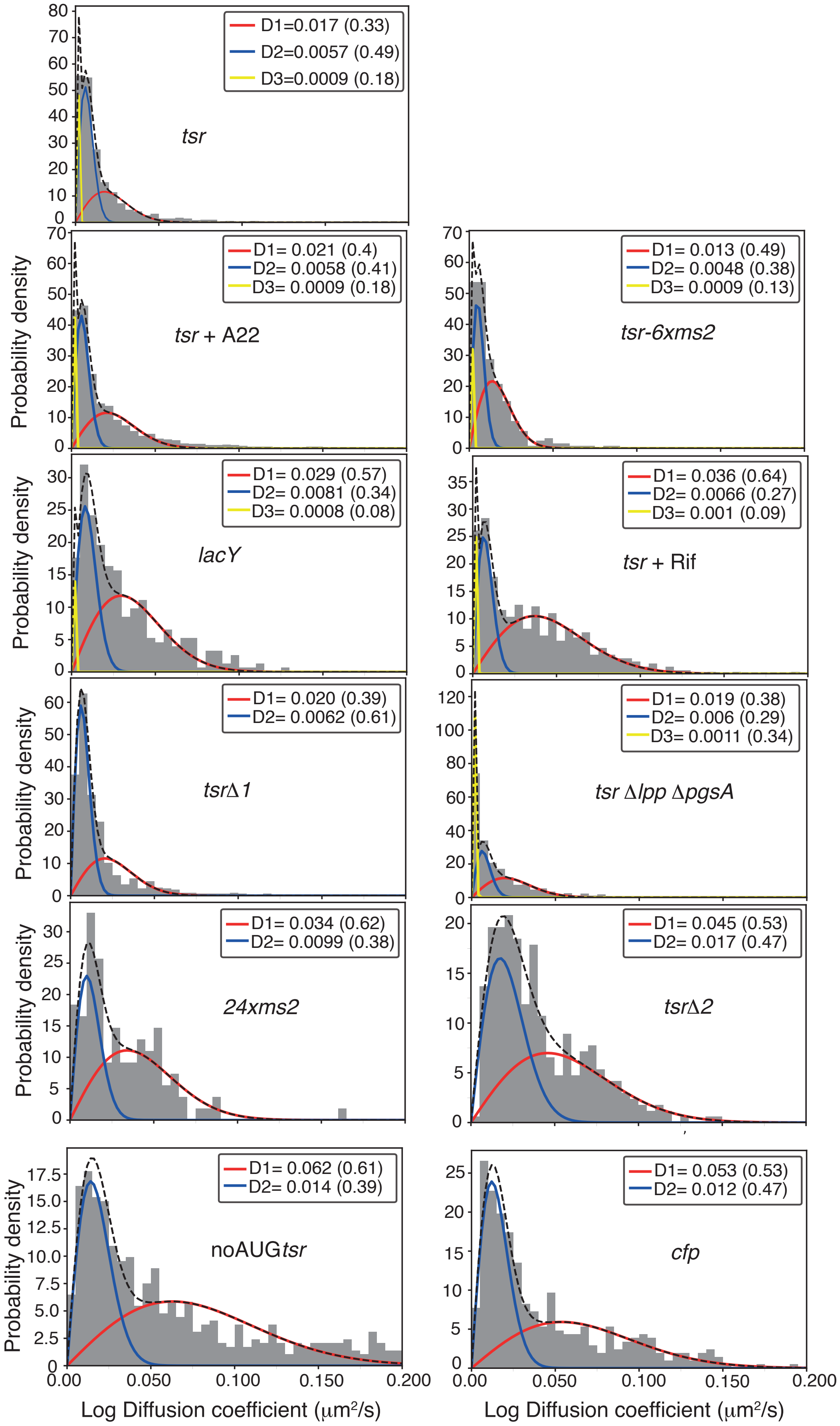

**Supplementary Figure 2**

Single time step-derived diffusion coefficients distributions for all data shown in Figure 2 in the main text. mRNA type is highlighted within each histogram.

### Supplementary Figure 3

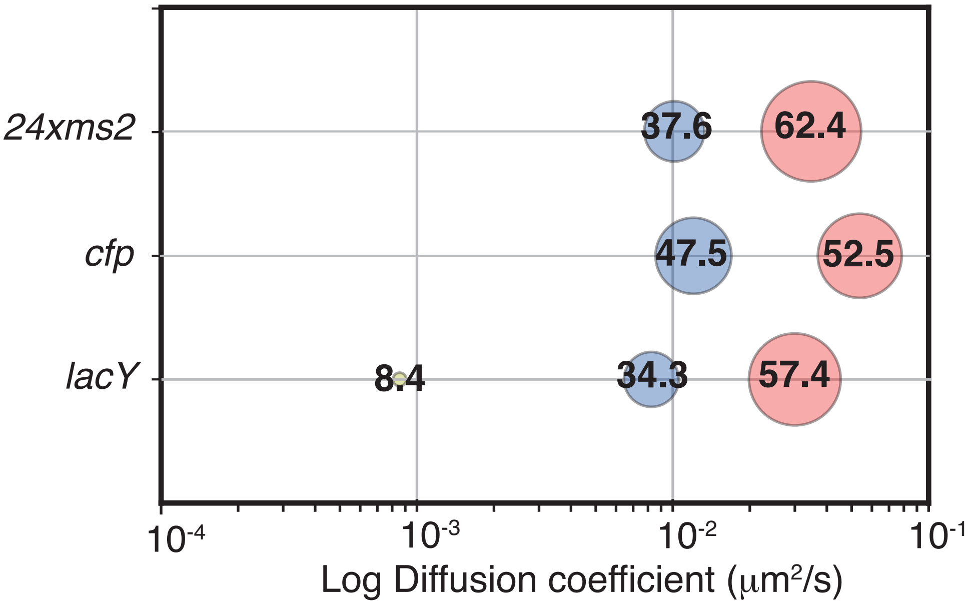


**Supplementary Figure 3**
Bubble plot showing the summary of the fitting procedure for different mRNA species. Numbers on top of bubbles and bubble size indicate the weight of each subpopulation. X-axis: log diffusion coefficient, $\mu m^2/s$. *24xms2* – mRNA molecule containing 24 MBS only and no coding mRNA. *cfp* – mRNA encoding cytoplasmic CFP protein, labelled with 24 MBS and expressed from a pZA21 plasmid. *lacY* – mRNA encoding membrane protein LacY, labelled with 24 MBS and expressed from a pZA21 plasmid.

### Supplementary Figure 4

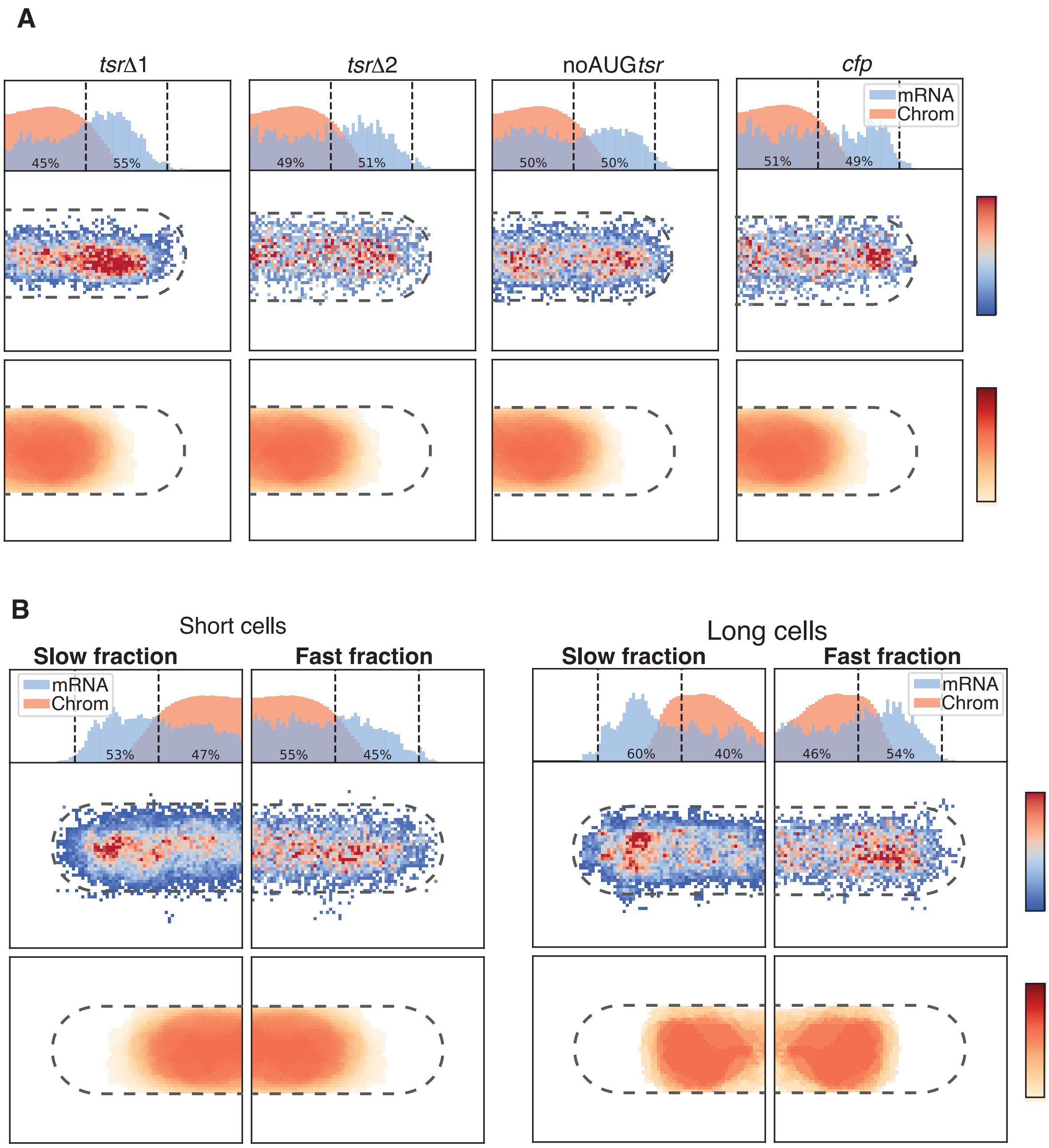


**Supplementary Figure 4**

a) Average localization maps of cells expressing *tsr*Δ1-*24xms2*, *tsr*Δ2-*24xms2*, noAUG*tsr*-*24xms2* and *cfp*-*24xms2* from pZA21 plasmid. Intermediate and fast subpopulations were binned. b) Average localization maps of cells expressing *tsr*-*24xms2* from pZA21 sorted into short and long cells. Average chromosome localization derived from experiments with pZA21-*tsr*-*24xms2* cells was plotted into folded up cells and mirrored to the right. On top, probability density along the x-axis of the cell of mRNA particles and the chromosome is shown. Numbers indicate the fraction of mRNA particles near the cell pole and cell center. Slow subpopulations represent binned slow and intermediate populations.
